# Natural variation in the roles of *C. elegans* autophagy components during microsporidia infection

**DOI:** 10.1101/418749

**Authors:** Keir M. Balla, Vladimir Lažetić, Emily Troemel

## Abstract

Natural genetic variation can determine the outcome of an infection, and often reflects the co-evolutionary battle between hosts and pathogens. We previously found that a natural variant of the nematode *Caenorhabditis elegans* from Hawaii (HW) has increased resistance against natural microsporidian pathogens in the *Nematocida* genus, when compared to the standard laboratory strain of N2. In particular, HW animals can clear infection, while N2 animals cannot. In addition, HW animals have lower levels of intracellular colonization of *Nematocida* compared to N2. Here we investigate how this natural variation in resistance relates to autophagy. We found that there is much better targeting of autophagy-related machinery to parasites under conditions where they are cleared. In particular, ubiquitin targeting to *Nematocida* cells correlates very well with their subsequent clearance in terms of timing, host strain and age, as well as *Nematocida* species. Furthermore, clearance correlates with targeting of the LGG-2/LC3 autophagy protein to parasite cells, with HW animals having much more efficient targeting of LGG-2 to parasite cells than N2 animals. Surprisingly, however, we found that *lgg-2* is not required to clear infection. Instead we found that loss of *lgg-2* leads to increased intracellular colonization in the HW background, although interestingly, it does not affect colonization in the N2 background. Altogether our results suggest that there is natural genetic variation in an *lgg-2*-dependent process that regulates intracellular levels of microsporidia at a very early stage of infection prior to clearance.

## Introduction

Natural genetic variation underlies differences in susceptibility to infection and inflammation among individuals.^1^ Genome-wide association studies in humans have revealed genetic variation in innate immune genes that predispose individuals to increased risk of infection and autoimmune disease. For example, polymorphisms in autophagy genes Nod2, ATG16L, and IRGM are associated with increased risk for Crohn’s disease, which is an inflammatory bowel disease characterized by a dysregulated gut microbiome.^2, 3^ The discovery that natural variation in autophagy genes can lead to differences in inflammation is part of a large body of evidence indicating a close connection between inflammation, intestinal immunity, and autophagy.^4, 5^

Autophagy is a broadly conserved cellular process that was originally studied for its role in degrading and recycling bulk intracellular material, and subsequently has been shown to play a role in immunity.^6^ When autophagy is used to degrade intracellular pathogens it is called xenophagy.^7-10^ Xenophagy often begins with localization of host ubiquitin to intracellular microbes, followed by localization of autophagy proteins, formation of an autophagosome to enclose the microbe, and then fusion with a lysosome to degrade microbial cargo. Initial studies of the role for autophagy in immunity focused on xenophagy, although it is now appreciated there are ‘non-canonical’ forms of autophagy that also promote anti-microbial defense.^11^ For example, autophagy can regulate secretion of anti-microbials during stress.^12^ Given that natural variation in autophagy underlies human risk of intestinal inflammatory disease, studying how autophagy responses vary among individuals in a species and how these responses contribute to immunity could provide insights into human disease.

The nematode *Caenorhabditis elegans* provides an attractive system for studying natural variation in innate immunity and the role of autophagy.^13^ Autophagy has been studied extensively in *C. elegans*.^14, 15^ Furthermore, there are many natural isolates of *C. elegans*, and various natural pathogens associated with these strains.^16^ The most common cause of infection for *C. elegans* in the wild is microsporidia, which comprise a phylum of >1400 species of obligate intracellular microbes related to fungi.^17, 18^ Microsporidia invade host cells using a polar tube that ‘fires’ to deliver a parasite cell called a sporoplasm directly into the host cell. The first microsporidian species shown to infect *C. elegans* was isolated from wild-caught nematodes near Paris, France and we named it *Nematocida parisii* strain ERTm1 (nematode-killer from Paris), because it causes a lethal intestinal infection.^19^ In the early stages of infection, *N. parisii* grows in direct contact with the host cytoplasm, and RNAseq analysis identified several predicted host ubiquitin ligase components induced by *N. parisii* infection.^20, 21^ Because ubiquitin ligases can act upstream of autophagy by targeting ubiquitin to microbes exposed to the cytoplasm,^22^ we explored the role of autophagy/xenophagy in resistance to *N. parisii*. *C. elegans* has conserved autophagy machinery, and other studies have demonstrated a role for autophagy in response to infection with extracellular, clinically relevant pathogens.^13, 23-26^ However, bona fide xenophagy (i.e. autophagy targeting to microbes that mediates subsequent clearance of pathogen) has not yet been demonstrated in *C. elegans*. We previously found that inhibiting autophagy by RNAi caused an increase in *N. parisii* load in the standard N2 laboratory strain of *C. elegans* from England, but only to a modest extent. Subsequently we have performed time course studies, and found that control animals are not able to clear infection.^21, 27^ Thus, we do not find evidence for xenophagic clearance of pathogens in the N2 host strain.

In contrast to N2, we found that a natural variant of *C. elegans* from Hawaii (hereafter referred to as HW) does have the ability to clear microsporidia from its intestinal cells.^27^ HW animals at the first larval (L1) stage can clear *Nematocida* infection, although they lose this ability later in life. In these previous studies we used a *Nematocida* strain (ERTm5) from Hawaii that we called *N. parisii* based on ribosomal sequence. We have subsequently performed whole genome sequencing and have described ERTm5 as a new species because it has only an average 85% amino acid identity with *N. parisii*; we named this new species *Nematocida ironsii*.^28^ Interestingly, the ability of HW to clear *N. ironsii* infection correlates with the developmental stage at which infection has fitness consequences for the host. In addition, HW animals exhibit lower levels of *N. ironsii* colonization early during infection, again only during the L1 stage. Quantitative genetic analysis indicated that the genomic regions that confer increased immunity contain genes predicted to encode ubiquitin ligase adaptor proteins.^27^ Therefore, we hypothesized that HW *C. elegans* may be better than N2 at targeting *Nematocida* cells with ubiquitin to direct autophagy machinery for clearance of these parasites via xenophagy. Indeed, because *C. elegans* lacks professional immune cells, and its intestinal epithelial cells are non-renewable, xenophagy provides an attractive candidate for a process that could explain how *C. elegans* can clear microsporidia infection from their intestinal cells.^29, 30^

Here we identify natural variation in both *C. elegans* and *Nematocida* in the targeting of host autophagy-related machinery to pathogen cells as well as the importance of autophagy-related machinery for infection levels. We found a striking correlation between targeting of autophagy-related machinery to *Nematocida* cells and the ability to clear these pathogens from host intestinal cells. For example, we found that HW animals have much higher levels of ubiquitin targeting to *Nematocida* cells compared to N2 animals. Furthermore, these differences are specific to the developmental stage at which HW animals can clear infection. We also observed higher frequencies of ubiquitin targeting around cells of a *Nematocida* species that is cleared, compared to two *Nematocida* species that are not cleared. In addition to ubiquitin, we observed higher frequencies of the autophagy protein LGG-2/LC3 targeted to *Nematocida* cells in HW animals compared to N2 animals. Surprisingly however, we found that LGG-2 is not required for clearance in HW animals, indicating that there are other proteins responsible for clearing this co-evolved pathogen. Instead, we found that LGG-2/LC3 regulates pathogen colonization. Interestingly, LGG-2/LC3 regulates intracellular colonization only in the HW strain, and not in N2. Previous work mapped HW resistance to genomic regions other than the region containing *lgg-2*^27^, and the LGG-2 protein does not appear to vary in amino acid sequence between N2 and HW (www.wormbase.org). Therefore, the *lgg-2* gene itself does not appear to underlie the genetic variation in pathogen resistance between N2 and HW. Rather, these findings indicate that an *lgg-2-*dependent process appears to naturally vary between N2 and HW, and is required to control levels of *Nematocida* parasites at a very early stage of infection.

## Materials & Methods

### *C. elegans* strains and maintenance

*C. elegans* were maintained on nematode growth medium seeded with *Escherichia coli* (OP50) as previously described.^31^ The N2 and CB4856 (HW in the text) strains were obtained from the CGC. All transgenic strains expressing fluorescent proteins carry extrachromosomal arrays. The N2 strain expressing GFP-tagged UBQ-1 (strain ERT261) was generated by injection of the pET341 plasmid *[vha-6p::GFP::ubq-1::unc-54 3’UTR; cb-unc-119(+)]* as previously described.^21^ CB4856 animals were injected with pET341 to generate HW GFP::UBQ-1 animals (strain ERT337). To generate transgenic mCherry::GFP-tagged LGG-1 animals we injected N2 or CB4856 animals with the pMH878 plasmid from the Hansen lab *[lgg-1p::mCherry::GFP::lgg-1::lgg-1 3’UTR]*.^32^ This yielded strains ERT426 and ERT429 in the CB4856 and N2 backgrounds, respectively. To generate transgenic GFP-tagged LGG-2 animals we injected QX1015 animals (*unc-119* mutant in CB4856 background) and bombarded EG6699 animals (*unc-119* mutant in N2 background) with the pRD117 plasmid from the Legouis lab *[lgg-2p::GFP::lgg-2::lgg-2 3‘UTR; cb-unc-119(+)]*.^33, 34^ This yielded strains ERT496 and ERT511, respectively. Information about all strains used in this study are presented in Table S1.

### *Nematocida* infection assays and imaging

Infection experiments were performed with *N. ironsii* strain ERTm5, *N. parisii* strain ERTm1, and *N. ausubeli* strain ERTm2.^17, 28, 35^ Spores were prepared and quantified as previously described.^36^ Host animals were either continuously exposed to *Nematocida* spores or pulse-inoculated with spores for 3 hours on plates as described in the figure legends. For Figure S7, animals were incubated in liquid culture for 15 minutes. Infections of L1 stage animals or L4 stage animals were initiated as described previously.^27^ Experiments were carried out at 20°C (Figures 1-3, S1B) or 25°C (Figures 4, 5, S1A, S4-S8). Animals were fixed at various times post-inoculation with 4% paraformaldehyde in PBS with 0.1% Tween 20 (PBST) for 45 minutes. N2, HW, GFP::UBQ-1 and GFP::LGG-2 transgenic animals were stained with the MicroB FISH probe conjugated to the CAL Fluor Red 610 dye at 5ng/μl overnight, then resuspended in PBST (clearance experiments) or Vectashield with DAPI (Vector Labs) to stain DNA (colocalization experiments). mCherry::GFP::LGG-1 animals were stained with the MicroB FISH probe conjugated to the Pacific Blue dye at 10ng/μl overnight, with no staining of DNA. For infection clearance experiments, the percentage of infected animals out of 100 total animals per sample was calculated as described previously.^27^ To image the localization of transgenic proteins during *Nematocida* infections, fixed and stained animals were mounted on 5% agarose pads and imaged using a 40X oil immersion objective on a Zeiss LSM700 confocal microscope run by ZEN2010 software. At least 10 infected transgenic animals were imaged per sample and *Nematocida* cells were counted as having transgenic proteins localized around them or not.

**Figure 1.**
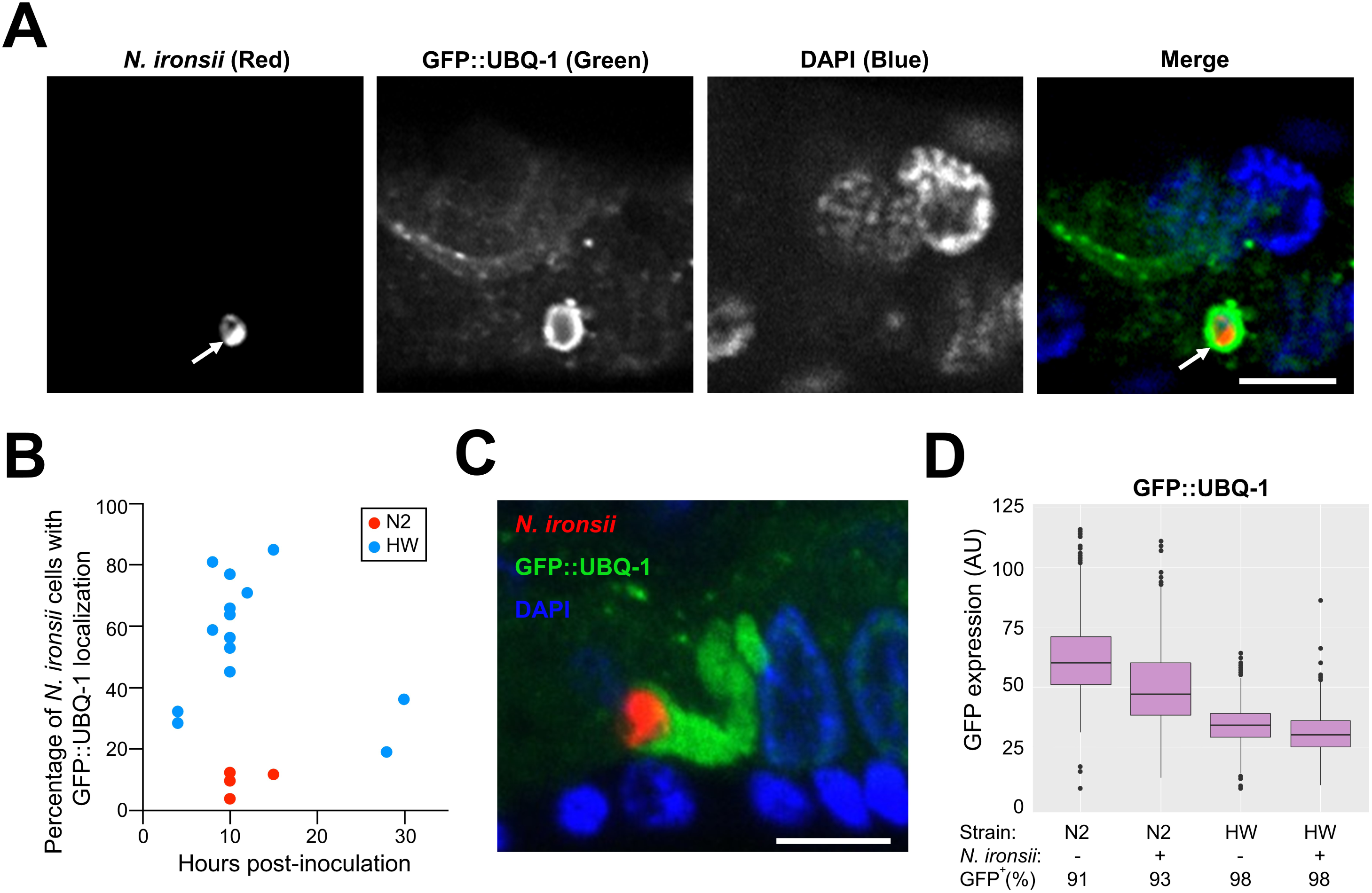
Host ubiquitin protein UBQ-1 localizes around a greater percentage of *N. ironsii* cells in HW animals than in N2 animals. (A) Example of UBQ-1 protein localized around *N. ironsii* cell in HW animal, fixed 10 hpi. *N. ironsii* cell (red FISH staining for rRNA) is shown in the first image, GFP-tagged transgenic host UBQ-1 protein (green) is shown in the second image, DNA stained with DAPI (blue) is shown in the third image, and a merged overlay is shown in the fourth image. The arrow points to an *N. ironsii* cell. Scale bar = 5 µm. (B) Percentage of *N. ironsii* cells that have UBQ-1 localization at different timepoints. Each dot indicates the percentage of UBQ-1 localization around all *N. ironsii* cells in an experiment. Blue dots represent the frequencies observed in HW animals; red dots represent frequencies in N2 animals. In 14 independent experiments, a total of 260 HW animals, and 406 *N. ironsii* cells were analyzed, while in 4 independent experiments, a total of 127 N2 animals and 310 *Nematocida* cells were analyzed. Each experiment included analysis of at least 10 animals, and each animal was infected by one to nine parasite cells. Data shown are compiled from more than three independent experiments per strain. (C) Abundant GFP::UBQ-1 (green) accumulation in close proximity to *N. ironsii* cell (red). Scale bar = 5 µm. From 4 independent experiments totaling 117 N2 animals with 296 *Nematocida* cells and 169 HW animals with 234 *Nematocida* cells, extreme amounts of ubiquitin targeting were observed in 10 out of 234 cells in HW, and were never observed in N2 animals. (D) Analysis of GFP::UBQ-1 expression in N2 and HW strains using a COPAS Biosrt machine, with expression measured on an individual animal basis. AU, arbitrary units. Boxplots show the interquartile range (IQR) from 25th to 75th percentile with horizontal lines indicating medians. The range bars encompass all data within 1.5 IQR above and below the upper and lower IQRs, respectively. Dots show individuals outside of that range. Only GFP levels in GFP^+^ animals are plotted, where GFP^+^ includes all animals with GFP levels above non-transgenic N2 animals. More than 1000 animals were measured per strain and condition.

### Measurements of transgenic protein levels

For Western blot measurements of protein levels, animals were washed off NGM plates with M9 buffer, and then washed once with PBS with 0.1% Tween 20 to remove bacteria. Samples were then resuspended in sample buffer with 1% SDS and 50 mM DTT and boiled at 95°C for 10 minutes. Lysates were then run on a 4-20% gradient SDS-PAGE gel (Bio-Rad) and transferred to PVDF membrane (Bio-Rad). The blots were stained with Ponseau S stain (Sigma-Aldrich) and imaged for total protein before staining with an anti-GFP antibody made in rabbits (a gift from Arshad Desai and Karen Oegema labs at UC San Diego) diluted at 1:5000 overnight at 4°C, then staining with an antirabbit HRP antibody at 1:10000 for 45 minutes at room temperature. The blots were treated with ECL reagent (Amersham GE Healthcare Life Sciences) and imaged on a Bio-Rad ChemiDoc. For measurements of GFP protein levels in live animals, uninfected and *N. ironsii* infected animals were collected 15 hour post-inoculation (hpi) and measured on a COPAS Biosort machine (Union Biometrica) for size (time-of-flight, TOF) and GFP (green). More than 1000 animals were measured per sample. Animals expressing transgenic GFP were distinguished from non-expressing animals by measuring GFP levels in wild type N2 animals and only analyzing GFP levels in animals above this threshold.

### CRISPR-Cas9 generation of *lgg-2* deletion alleles

Using the CRISPR design tool (http://crispr.mit.edu), DNA sequences with high scores were chosen in regions upstream from the start and downstream from the stop codon of *lgg-2* to create crRNAs (synthesized by Integrated DNA Technologies). crRNAs were annealed to the tracrRNA, and sgRNA products were co-injected with Cas9 protein and *dpy-10* sgRNA marker.^37^ Dumpy F1 animals were screened for *lgg-2* deletions using PCR analysis. Isolated homozygous lines were confirmed and characterized by DNA sequence analysis and backcrossed three times to the original strain, CB4856 for *jy44* and *jy45* alleles, and N2 for *jy102* and *jy103*. The *jy44* allele contains a 1376-nucleotide deletion, which removes the entire *lgg-2* gene and an additional 132 nucleotides upstream and 146 nucleotides downstream of the gene. Similarly, the whole *lgg-2* gene together with 138 nucleotides upstream and 146 nucleotides downstream (1382 nucleotides in total) are substituted for a single nucleotide insertion in *jy45* allele. In the N2 background, the *jy102* allele contains a 1356-nucleotide deletion, which removes the whole *lgg-2* gene together with 125 nucleotides upstream and 133 nucleotides downstream of the gene; the *jy103* allele is an indel with 7 nucleotides inserted in place of 1344 nucleotides, which spans the entire *lgg-2* gene together with 122 nucleotides upstream and 124 nucleotides downstream of the gene.

### Sequence of *lgg-2* in N2 and CB4856

According to www.wormbase.org, there are no predicted amino acid changes in LGG-2 between N2 and CB4856. There is one nucleotide substitution in *lgg-2* in CB4856 animals compared to N2 animals (WBVar00191911): A/T in the first intron. Also, there is one nucleotide substitution in a possible promoter region (WBVar00191912): G/A.

### Gene expression measurements

To confirm the absence of *lgg-2* mRNA transcripts in *lgg-2(jy44)* mutants, synchronized populations of CB4856 wt and HW *lgg-2(jy44)* animals were grown at 20°C to reach L4 stage, and then collected in TriReagent (Molecular Research Center, Inc.), and RNA extraction was performed following manufacturer’s guidelines. cDNA synthesis was performed using SuperScript™ VILO™ cDNA Synthesis Kit (Thermo Fisher Scientific) following manufacturer’s guidelines. qRT-PCR was performed as using a BioRad CFX Connect. *lgg-2* mRNA was amplified in two separate reactions using the same forward primer (5’-GAATCGTTCCATCGTTCAAGG-3’) and two different reverse primers (5’-TGTTGGCTGCGGATTTCT-3’ and 5-TTGGAGGCGTCGTCTAACA). Each of two replicate experiments was measured in duplicate and compared to the expression of *snb-1* control gene.

### Feeding rate measurements

Fluorescent beads (Fluoresbrite™ Polychromatic Red 0.5 µm Microspheres, Polysciences, Inc.) were added in a 1:50 ratio to a mix of starved L1 animals, concentrated OP50 and *N. ironsii* spores in M9 buffer. This mix was prepared following the protocol for clearance experiments. After 30 minutes incubation at 25°C, animals were fixed in 4% paraformaldehyde and analyzed. For Figure S5A, animals were imaged using a Zeiss AxioImager; fluorescence levels were analyzed using the FIJI program. Mean fluorescence per animal was calculated for 50 animals per strain, per replicate, in three separate experiments. Background fluorescence levels were subtracted from obtained results. For Figures S5B and S8B, bead fluorescence was analyzed using COPAS Biosort machine and standardized to TOF. Based on the fluorescence intensity of beads in a given experiment, Photomultiplier Tube (PMT) settings were adjusted accordingly. For experiments shown in Figure S5B the PMT setting was 800, while for Figure S8B it was 700. Based on the flow rate in a given experiment, TOF cutoffs are adjusted accordingly. For Figure S5B the TOF cutoff was 70-200, and for Figure S8B it was 100-200.

## Results

### *N. ironsii* cells are surrounded by ubiquitin more frequently inside the Hawaiian strain of *C. elegans* than the N2 strain

To investigate the hypothesis that HW *C. elegans* clear *N. ironsii* infections through improved targeting of ubiquitin to parasite cells followed by autophagic clearance, we examined the localization of ubiquitin to parasite cells in the HW host compared to the N2 host strain, which does not clear infections. First, we generated transgenic N2 and HW strains that express GFP-tagged ubiquitin under the control of the intestinal-specific promoter *vha-6* (GFP::UBQ-1). We pulse-infected these GFP::UBQ-1-expressing strains at the first larval (L1) stage by feeding animals with *N. ironsii* spores for three hours, then removed them from spores and fixed them at different hours post-inoculation (hpi). We performed FISH staining on these fixed animals using a probe that targets *Nematocida* rRNA to label parasite cells and then determined the percentage of those cells that also have GFP::UBQ-1 localization (Figure 1A, B). With this method, we found that a much higher percentage of parasite cells had ubiquitin localization to parasite cells in HW animals than in N2 animals. For example in one experiment at 15 hpi, we found that 84% of *Nematocida* cells (out of 19 total parasite cells analyzed in 10 separate animals) were targeted with GFP::UBQ-1 in HW animals (Figure 1B), and every animal had at least one parasite cell with GFP::UBQ-1 localization. In contrast, only 11% of *Nematocida* cells (out of 37 total parasite cells analyzed in 10 separate animals) were targeted with GFP::UBQ-1 in N2 animals at 15 hpi (Figure 1B). Interestingly, in some HW animals we observed exuberant accumulations of ubiquitin that formed long structures next to parasite cells, which were never observed in N2 animals (Figure 1C).

Next we analyzed how ubiquitin targeting to parasites correlates with their clearance. In previous clearance experiments at 25°C we had found parasite load much lower in HW animals at 20 hpi compared to 3 hpi, but did not measure when clearance began between those two timepoints.^27^ Here we analyzed clearance at several timepoints after 3 hpi and saw that clearance begins around 10 hpi at 25°C (Figure S1A). Because we have found that there are more protein aggregates at 25°C, which can make localization experiments challenging,^21^ we performed our autophagy localization experiments at 20°C (this paper and ^21^). (We do note that previous studies indicate that autophagy does not appear to proceed differently at 20°C compared to 25°C).^32^ Therefore, we also analyzed clearance at 20°C to ensure that clearance is not just a high-temperature phenomenon, and to better compare the kinetics of clearance and ubiquitin targeting. We found that parasite clearance also occurs at 20°C (Figure S1B). Furthermore, we were able to compare the kinetics of clearance with ubiquitin targeting at the same temperature, and found that the peak of ubiquitin targeting correlates with the start of clearance (compare Figure 1B and Figure S1B). These findings are consistent with ubiquitin targeting driving parasite clearance.

One potential explanation for increased targeting of ubiquitin to parasite cells in HW animals is increased expression from the GFP::UBQ-1 transgene in HW animals. To investigate this possibility, we compared protein levels of GFP::UBQ-1 with two methods. First, we performed Western blots with anti-GFP antibodies on lysates from transgenic HW and N2 animals to compare levels of GFP::UBQ-1 across populations of animals. Here we found that HW animals do not have increased levels of GFP::UBQ-1 compared to N2 animals (Figure S2). Second, we analyzed GFP::UBQ-1 fluorescence on a per-animal level using a COPAS Biosort ‘worm sorter’. While there was animal-to-animal variability in fluorescence levels of GFP-ubiquitin, the overall levels were not higher in HW compared to N2 animals (Figure 1D). Therefore, the increased targeting of GFP-ubiquitin to *N. ironsii* cells in HW compared to N2 does not appear to be simply due to increased GFP::UBQ-1 levels in HW animals.

### Localization of ubiquitin to parasite cells correlates with their clearance in terms of host age and parasite species

In previous work, we showed that HW animals can clear infection by *N. ironsii* only during the L1 larval stage, and that this ability is lost in later larval stages.^27^ To characterize genetic variation in clearance on the parasite side, we infected HW at the L1 and L4 stages with three different *Nematocida* species. We observed variation in clearance of these *Nematocida* species by HW animals, where *N. ironsii* was cleared by L1, but not L4 animals while *N. parisii* and *N. ausubeli* were cleared poorly by both L1 and L4 animals (Figure 2A, B). Therefore, the clearance of microsporidia by HW animals is specific to *N. ironsii* infections at the L1 stage.

**Figure 2.**
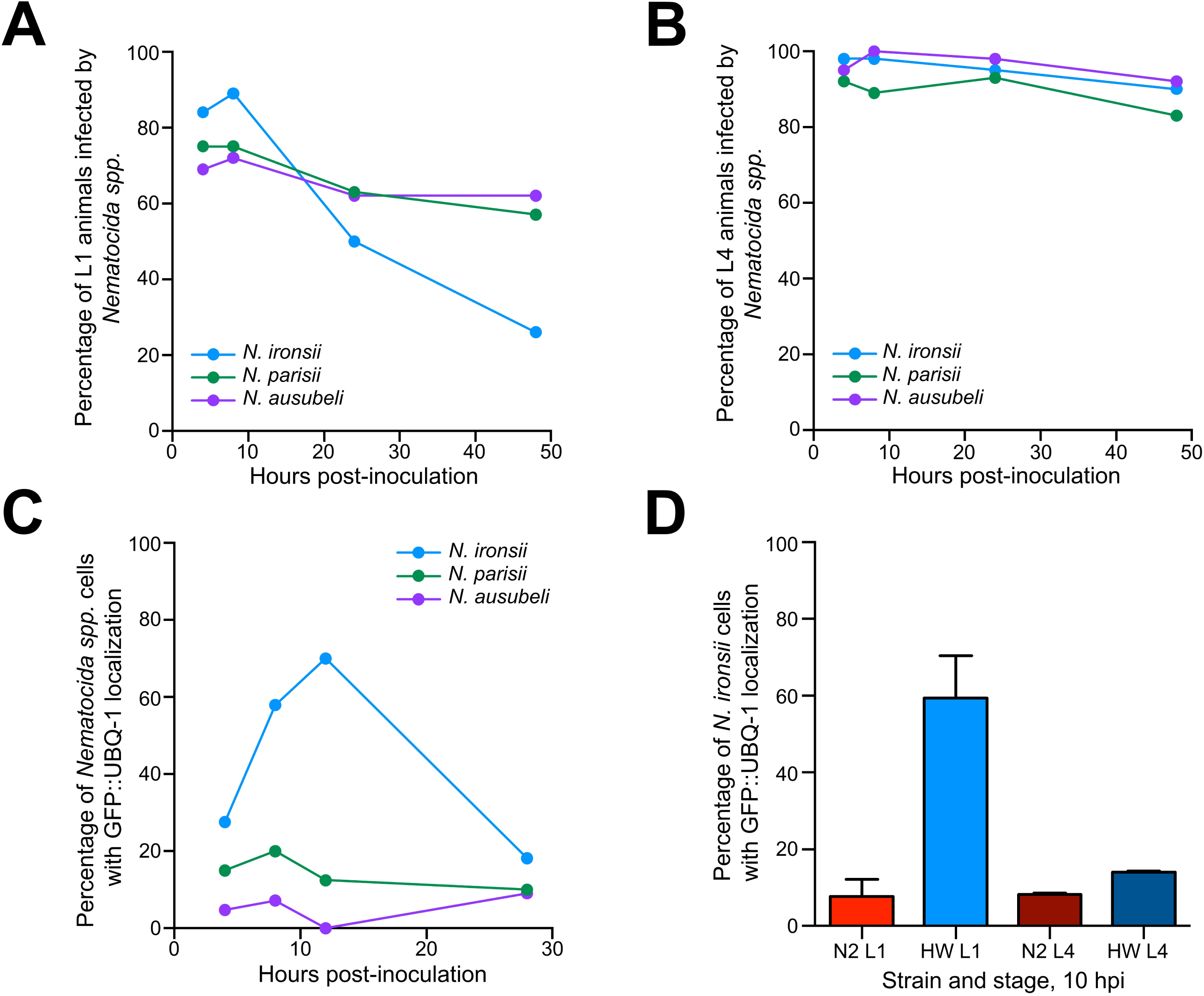
HW ability to clear microsporidia infection depends on host developmental stage and microsporidia species. (A-C) Each dot represents an experiment. (A) HW L1 animals are more efficient in clearing *N. ironsii* infection (blue line) than *N. parisii* (green line) or *N. ausubeli* (purple line) infections. Animals were pulse-inoculated with spores of different *Nematocida* species, and the percent of infected animals (y-axis) was analyzed at different time points (x-axis). (B) HW animals infected at the L4 stage, using the same experimental setup as in (A), are unable to eliminate *Nematocida spp*. (C, D) The localization of host ubiquitin around *Nematocida spp.* cells is most frequent in contexts where infection is eliminated. (C) Transgenic GFP::UBQ-1 HW L1 animals were pulse-inoculated with *Nematocida spp.* spores and fractions of the population were fixed at several time points afterwards to assess the frequency of UBQ-1 localization around *N. ironsii* (blue line), *N. parisii* (green line), or *N. ausubeli* (purple line) cells. Data shown are representative of two independent experiments. (D) Transgenic GFP::UBQ-1 N2 and HW animals were inoculated with *N. ironsii* spores at the L1 stage or L4 stage and fixed 10 hpi to quantify the frequency of UBQ-1 localization around *N. ironsii* cells. Data for L1 infections are from six independent experiments, data for L4 infections are from two independent experiments. Averages are shown with standard deviations. (A-D) All experiments were carried out at 20°C.

To determine how these levels of clearance relate to the association of host ubiquitin with parasite cells, we examined the frequency of ubiquitin localization to *Nematocida* cells in L1 HW animals over time. Here we observed a high frequency of ubiquitin localization around *N. ironsii* cells but not *N. parisii* or *N. ausubeli* cells (Figure 2C). Furthermore, the high frequency of ubiquitin localization to *N. ironsii* cells was observed in HW animals infected at the L1 stage but not in HW animals infected at the L4 stage or N2 animals at either stage (Figure 2D). Thus, the frequency of host ubiquitin localization around microsporidia cells in *C. elegans* is highly correlated with clearance of infection.

### Localization of LGG-2/LC3 but not LGG-1/GABARAP correlates with parasite clearance

Xenophagic clearance of intracellular pathogens involves targeting of autophagic machinery after ubiquitin targeting to pathogen cells. One of the most well-studied autophagic events is activation and conjugation of proteins in the LC3/GABARAP family to the developing autophagosome, which are required for fusion with the lysosome.^38^ Therefore, we examined whether LGG-1 (homolog of GABARAP in mammals) or LGG-2 (homolog to LC3 in mammals), localized more frequently to parasite cells in HW worms compared to N2 worms. To analyze localization, we first generated N2 and HW transgenic animals with *lgg-1p::mCherry::GFP::LGG-1* or *lgg-2p::GFP::LGG-2* transgenes. Then we performed experiments similar to the ubiquitin studies in Figure 1. We pulse-infected these animals for 3 hours, fixed them at various timepoints after inoculation, stained with a FISH probe to label *N. ironsii* cells and then quantified parasite localization by following GFP (Figure 3A). Here we found that there was a gradual increase in mCherry::GFP::LGG-1 targeting to *N. ironsii* cells over time in both N2 and HW animals, with no obvious difference between the two strains (Figure 3B). In contrast, we found that there was a much greater percentage of parasite cells targeted with GFP::LGG-2 in HW animals, compared with N2 animals (Figure 3B). This increased localization of LGG-2 was comparable to the increased ubiquitin targeting seen in HW compared to N2 (Figure 1C).

**Figure 3.**
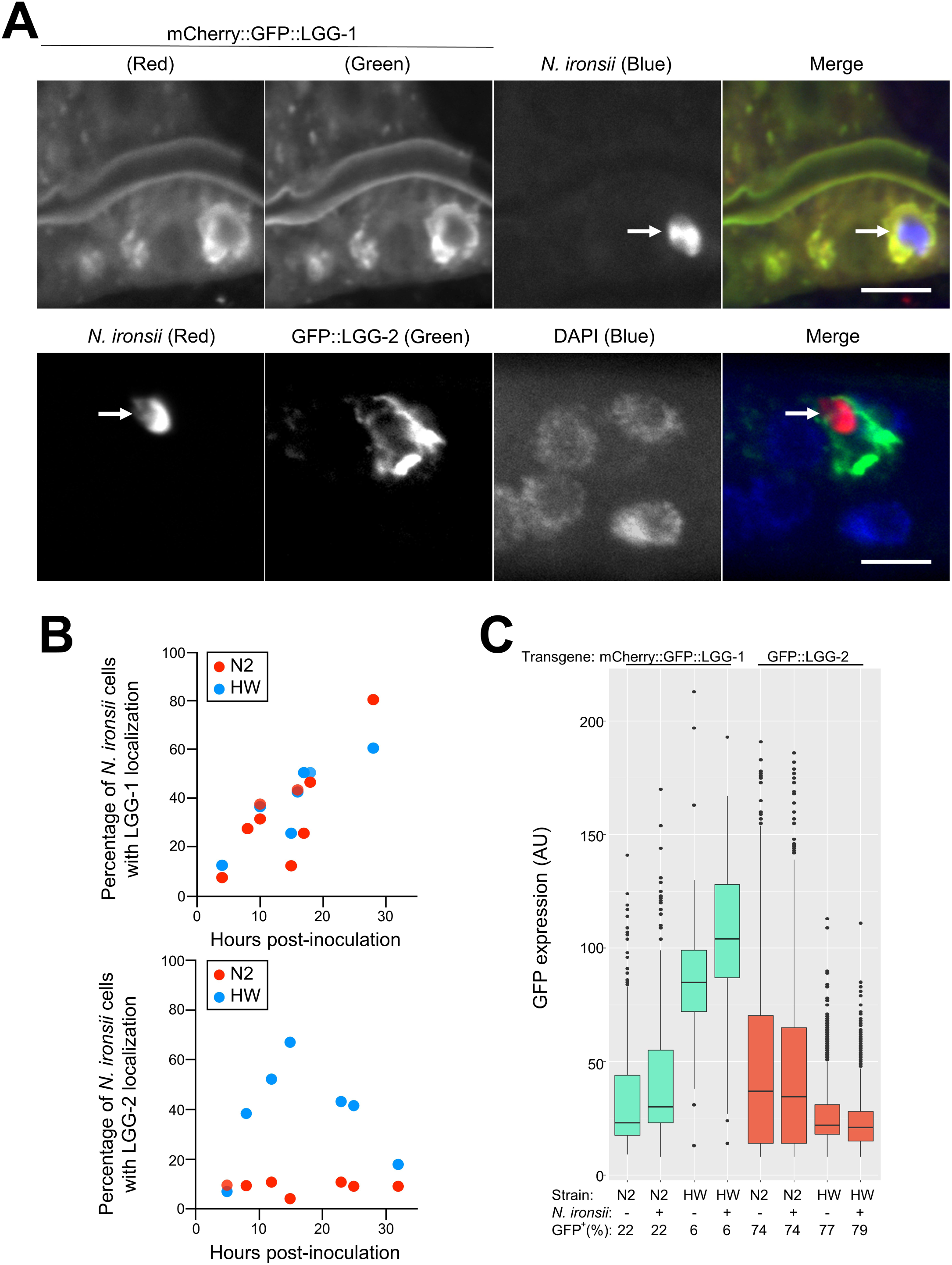
Localization of host autophagy proteins LGG-1 and LGG-2 to microsporidia cells in HW and N2 animals. (A) Examples of autophagy host proteins LGG-1 and LGG-2 localized around *N. ironsii* cells. The upper row shows mCherry-tagged transgenic host LGG-1 (red) in the first column, GFP-tagged transgenic host LGG-1 (green) in the second column, *N. ironsii* (blue) in the third column, and a merged image in the fourth column. The lower row shows *N. ironsii* FISH staining for rRNA (red) in the first column, GFP-tagged LGG-2 (green) in the second column, DNA stained with DAPI (blue) in the third column, and a merged image in the fourth column. Scale bars = 5 µm. The arrows point to *N. ironsii* cells. The images for mCherry::GFP::LGG-1 were from transgenic HW animals fixed 16 hpi and for GFP::LGG-2 12 hpi. (B) Percentage of *N. ironsii* cells with localization of host transgenic fusion proteins (LGG-1 in the upper graph, LGG-2 in the lower graph) over time. Each dot represents an experiment showing average frequency of localization around more than 10 *N. ironsii* cells per sample, with at least 10 animals per experiment. Blue dots represent the frequencies observed in HW animals; red dots represent frequencies in N2 animals. Data shown are compiled from more than three independent experiments per strain. (C) GFP-fusion autophagy protein analysis in transgenic N2 and HW strains. Transgenic GFP expression measured on an individual animal basis. AU, arbitrary units. As in Figure 1, boxplots show the interquartile range (IQR) from 25th to 75th percentile with horizontal lines indicating medians. The range bars encompass all data within 1.5 IQR above and below the upper and lower IQRs, respectively. Dots show individuals outside of that range. Only GFP levels in GFP^+^ animals are plotted, where GFP^+^ includes all animals with GFP levels above non-transgenic N2 animals. More than 1000 animals were measured per strain and condition.

To eliminate the possibility that the increased targeting of LGG-2 in HW animals was simply due to increased levels of GFP::LGG-2 expression, we performed Western blot analysis (Figure S3) and COPAS Biosort ‘worm sorter’ analysis of GFP::LGG-2 (and mCherry::GFP::LGG-1) (Figure 3C). Similar to our analysis of ubiquitin levels, we did not see an increased level of GFP::LGG-2 (or mCherry::GFP::LGG-1) expression in HW compared to N2 animals.

### LGG-2/LC3 is not required for clearance of *N. ironsii*

The increased localization of LGG-2 to *N. ironsii* cells in HW, and the ability of this strain to clear infection led us to hypothesize that parasite cells associated with LGG-2 were destined to be cleared, and that LGG-2 was important for the clearance. Therefore, we examined a functional role for LGG-2 in defense against *N. ironsii*. To do this we used CRISPR/Cas9 editing to generate a complete deletion of the *lgg-2* locus in the HW strain background. We isolated and characterized two independent alleles (called HW *lgg-2(jy44) and* HW *lgg-2(jy45)*) in which the entire *lgg-2* locus is deleted and should produce no LGG-2 protein (Figure 4A). We confirmed via qRT-PCR that *lgg-2* mRNA was not produced in *lgg-2(jy44)* mutant animals (Table S2). *lgg-2* mutant strains were backcrossed three times and then analyzed for their ability to clear *N. ironsii*. In several independent experiments, we saw that HW *lgg-2* mutants were still able to clear *N. ironsii* between 3 hpi and 20 hpi (Figure 4B, Figure S4A). Therefore, despite the striking increase in localization of LGG-2 to *N. ironsii* cells in HW animals compared to N2 animals, LGG-2 is not required for clearance of *N. ironsii* under these conditions.

**Figure 4.**
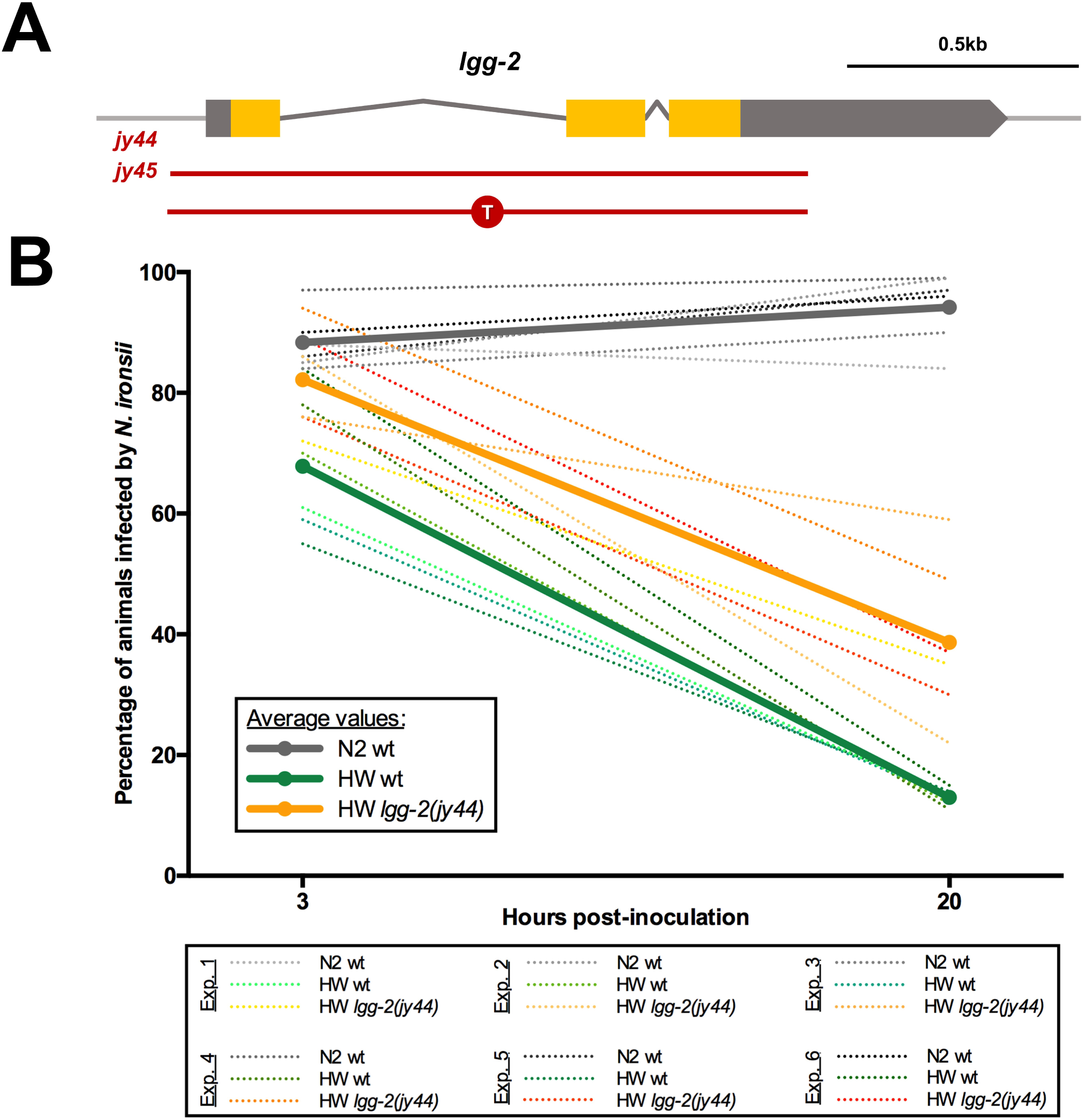
LGG-2 is not required for *N. ironsii* clearance. (A) Schematic representation of *lgg-2* gene. Orange boxes indicate exons, gray boxes indicate untranslated regions (UTRs). Red lines mark the deletion allele *jy44* and the indel allele *jy45* (inserted thymidine is indicated in the red circle). (B) *N. ironsii* clearance in N2 wild-type (wt) animals (gray lines), HW wild-type (wt) (green lines) and HW *lgg-2* mutant animals (orange lines). Thick lines represent average values of six experiments; dotted lines indicate results from individual experiments. 100 animals were analyzed per strain per time point (3 hpi and 20 hpi). Experiments were performed at 25°C.

### LGG-2 regulates *N. ironsii* colonization of intestinal cells in HW animals but not in N2

In addition to differences in clearance ability, our previous studies indicated that N2 animals have an increase in intracellular colonization at 3 hpi compared to HW animals, but only at the L1 stage.^27^ Consistent with this observation, we noticed in our clearance assays here that N2 animals had a higher percentage of infected animals than HW animals (Figure 4B). Interestingly, we also saw that HW *lgg-2* mutants had a higher percentage of animals infected with *N. ironsii* compared to HW wild-type animals, with HW *lgg-2* mutants having a percentage of infected animals closer to N2 wild-type animals (Figure 4B, S4A). When we analyzed pathogen load, we found that HW *lgg-2* animals had more sporoplasms per animal than HW wild-type animals at 3 hpi, with levels similar to N2 wild-type animals (Figure 5A, S4B). To ensure that the higher pathogen load phenotype was specific to *lgg-2(jy44)*, we crossed the GFP::LGG-2 construct used for localization into HW *lgg-2* mutant animals and found that this construct successfully rescued the higher pathogen load phenotype of HW *lgg-2(jy44)* mutants (Figure 5B).

**Figure 5.**
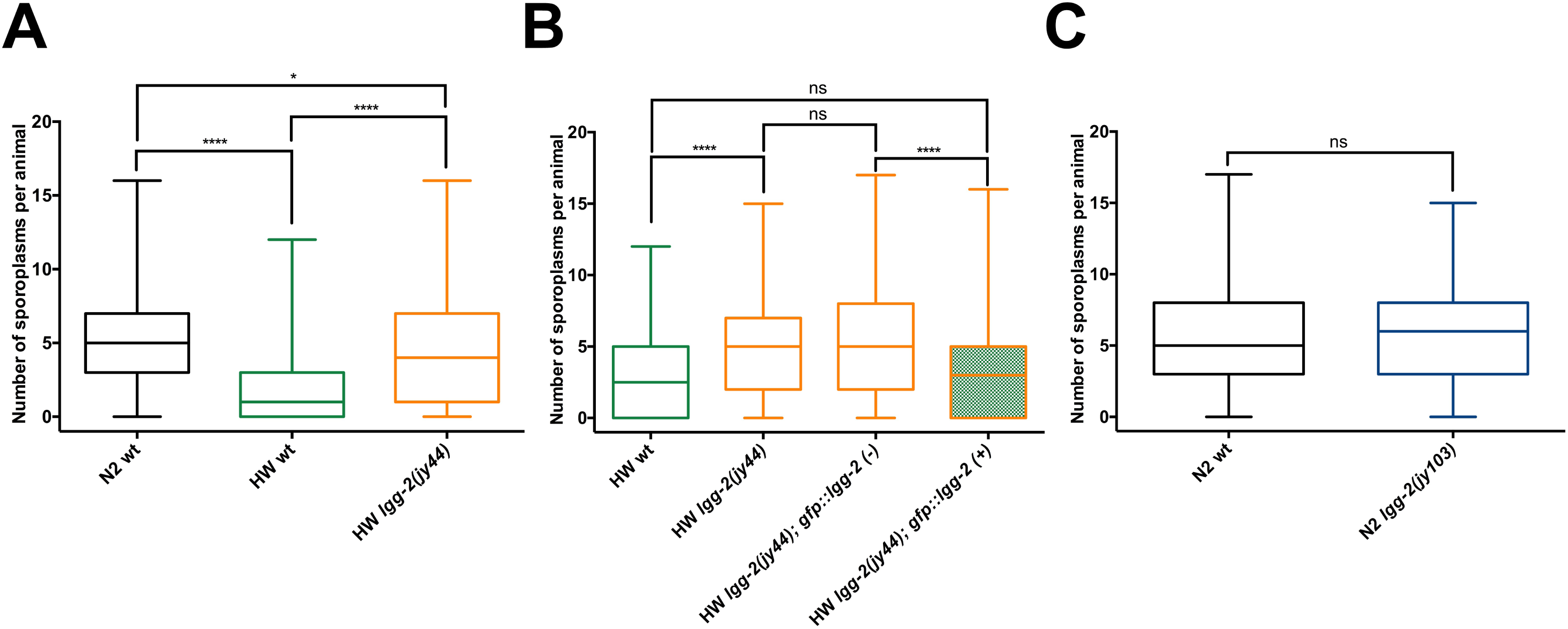
LGG-2 regulates intracellular colonization of *N. ironsii* in HW but not in N2 *C. elegans*. (A) HW *lgg-2* mutants on average have significantly higher *N. ironsii* colonization in comparison to HW wild-type animals, and similar infection rate to N2 animals, under the same experimental conditions. 600 animals were analyzed for each strain. (B) GFP::LGG-2 expression in *lgg-2(jy44)* mutants is able to rescue the higher pathogen load phenotype. 300 animals were analyzed for each genotype. (C) *lgg-2* deletion in N2 background does not affect *N. ironsii* colonization. 300 animals were analyzed for each strain. (A-C) Results from six (A) or three (B, C) independent experiments are shown as box-and-whisker plots, indicating the number of *N. ironsii* sporoplasms per animal at 3 hpi. Each box represents 50% of the data closest to the median value (line in the box). Whiskers span the values outside of the box. A student’s t-test was used to calculate *p* values; *p* < 0.001 is indicated with four asterisks; *p* < 0.05 is indicated with one asterisk; ns indicates nonsignificant difference (*p* > 0.05). Experiments were performed at 25°C.

One potentially trivial explanation for the higher pathogen load in HW *lgg-2* mutants could be that they have greater exposure to the pathogen, because it accumulates to a higher level within the intestinal lumen either through increased feeding or decreased defecation. To examine this possibility, we used a fluorescent bead-feeding assay and measured accumulation of beads inside the intestinal lumen in HW and HW *lgg-2* mutants. Here we found that there was no increase in the accumulation of beads in HW *lgg-2* mutants, indicating that their increased parasite load at 3hpi is likely not due to increased exposure to parasites (Figure S5). Therefore, *lgg-2* appears to be acting at a very early stage to regulate pathogen load of the natural intracellular parasite *N. ironsii* in HW animals.

Although clearance still occurred in HW *lgg-2* mutants, these animals had slightly less efficient clearance between 3 hpi and 20 hpi compared to HW wild-type animals (Table S3). To determine whether this was due to the difference in initial dose at 3 hpi between HW and HW *lgg-2* mutants, we infected HW animals with a higher spore dose than HW *lgg-2(jy44)* animals such that both strains had roughly equivalent initial sporoplasm number at 3 hpi (Figure S6). Under this condition, we found that clearance from 3 hpi to 20 hpi was similar in HW and HW *lgg-2* mutants, indicating that *lgg-2* is not required for clearance of *N. ironsii*, but is required for regulating the initial colonization of this natural parasite inside host intestinal cells.

We next considered the possibility that HW wild-type animals may already have cleared some infection before being analyzed for pathogen load at 3 hpi. If this were the case, then the increase in sporoplasm number in *lgg-2(jy44)* mutant animals at this timepoint would be a defect in clearance, rather than an increase in colonization. To test this hypothesis, we infected N2, HW wild-type and HW *lgg-2(jy44)* animals with a high dose of *N. ironsii* spores for 15 minutes and then analyzed the number of sporoplasms per animal. Interestingly, even after this very short-term incubation, HW *lgg-2(jy44)* mutants showed higher pathogen load than the HW wild type, suggesting that the intestinal cells in these mutants are more easily colonized by *N. ironsii* (Figure S7).

Because N2 wild-type animals have a higher initial colonization rate of *Nematocida* compared to HW wild-type animals, we hypothesized that some *lgg-2*-dependent process active in HW animals (that is inactive in N2) might be responsible for this difference. If this were the case, then loss of *lgg-2* in the N2 background would not have an effect. To test this hypothesis, we generated two *lgg-2* deletion alleles (*jy102* and *jy103*) in the N2 genetic background and tested for pathogen load at 3 hpi. Notably, we observed similar infection rates in wild-type and *lgg-2(jy103)* mutant animals (Figure 5C). We confirmed this result using the other deletion allele, *lgg-2(jy102)* (Figure S8A), and also showed that there is no significant difference in bead accumulation between N2 wild type and *lgg-2* mutant animals (Figure S8B). Therefore, the role of LGG-2 in pathogen colonization appears to be specific to the HW background, and the difference in colonization between N2 and HW appears to depend on LGG-2.

## Discussion

These studies reveal natural variation in the role of *C. elegans* autophagy-related machinery in response to natural *Nematocida* pathogens (Figure 6). We observe natural variation in localization of ubiquitin to parasite cells in different strains, and this localization correlates very well with clearance of parasite. In addition, there is a correlation between clearance and localization of the autophagy protein LGG-2/LC3 to *N. ironsii* cells, with HW animals having increased targeting compared to N2 animals. These findings led us to hypothesize that parasite cells associated with LGG-2/LC3 are cleared, which could explain the greater resistance of HW animals compared to N2. However, we found that HW *lgg-2* mutants can still clear infection, which does not support this hypothesis. Instead, we found that *lgg-2* regulates the level of microsporidia colonization inside intestinal cells in HW animals, but not in N2 animals. The LGG-2 predicted amino acid sequence does not vary between N2 and HW (see Materials and Methods). Furthermore, quantitative genetic studies indicate that the basis for the N2 and HW difference maps to chromosomes II, III and V, and not to chromosome IV, where *lgg-2* resides.^27^ Therefore, we do not favor a hypothesis that *lgg-2* itself is responsible for the genetic variation between these two strains. Instead, we propose that an *lgg-2*-dependent process (and not *lgg-2* itself) is active in HW animals to regulate levels of colonization, and is not active in N2 animals. In addition, our findings indicate that an *lgg-2*-independent process mediates clearance.

**Figure 6.**
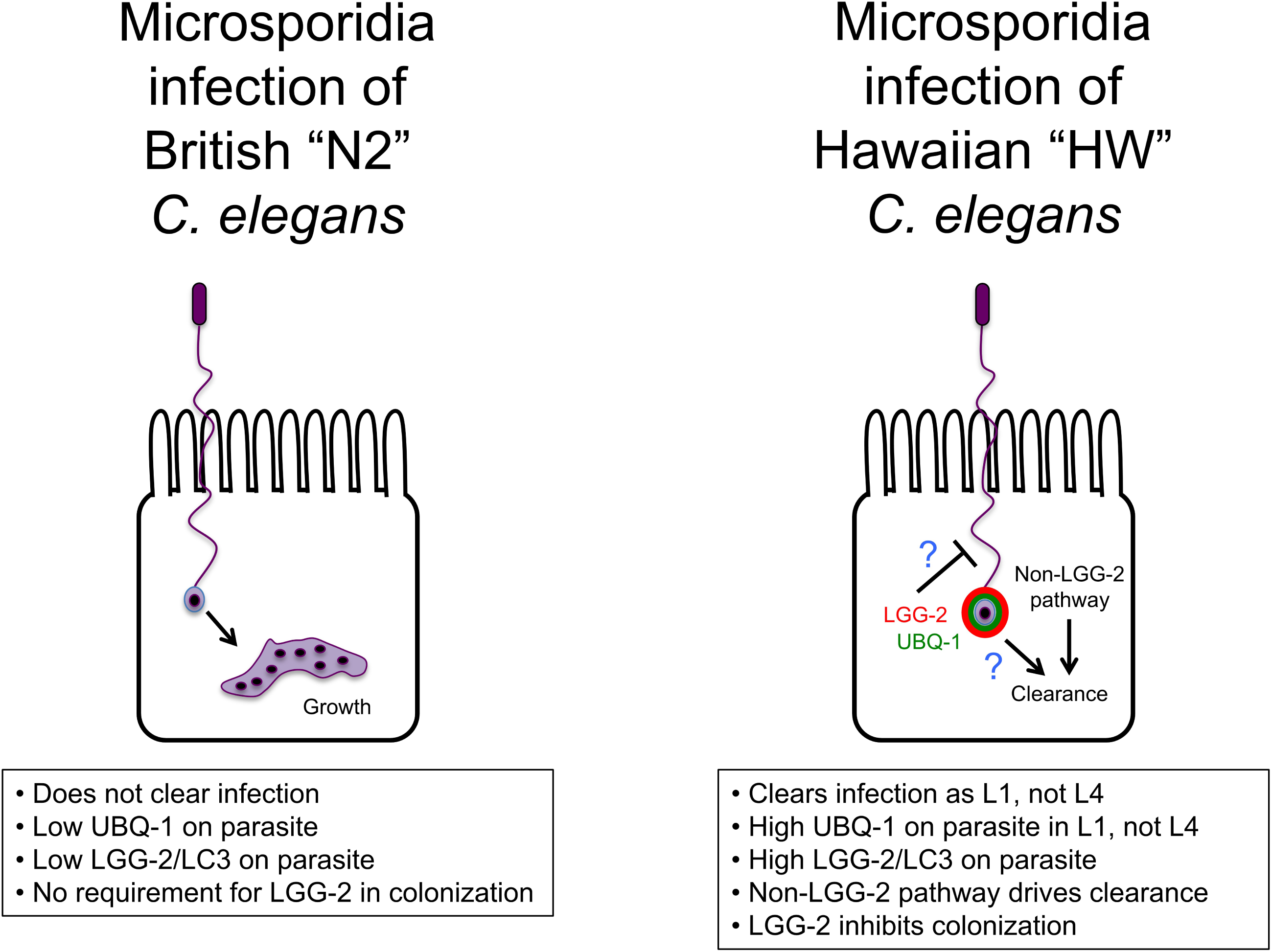
Model for invasion and clearance of *N. ironsii* in HW *C. elegans* hosts. N2 *C. elegans* fails to clear *N. ironsii* infection, while HW *C. elegans* clears *N. ironsii* as L1 animals, when there is high ubiquitin and LGG-2 targeting. An LGG-2-independent pathway mediates clearance, and LGG-2 inhibits initial colonization of *N. ironsii* inside host intestinal cells only in HW animals.

What non-LGG-2 protein mediates clearance of *Nematocida* cells in HW animals? In mammals there are six proteins in the LC3/GABARAP family that contribute redundantly to autophagy-mediated clearance.^38^ *C. elegans* has two proteins in this family, LGG-2 (homolog of LC3) and LGG-1 (homolog of GABARAP), and they do have redundant function in other contexts.^33, 34^ Therefore, it is possible that LGG-1 (GABARAP ortholog) is able to compensate for LGG-2 in clearance of microsporidia in HW *C. elegans*. It is difficult to test this model because LGG-1 is an essential gene, and HW worms are resistant to germline RNAi,^39, 40^ which is required to perform RNAi knockdown treatments that will affect L1 larvae where we see the resistance phenotype. However, we find it unlikely that LGG-1 could substitute for LGG-2 given the extensive LGG-1 localization to parasite cells we observed in both resistant HW and susceptible N2 animals (see Figure 3).

Our findings do not rule out LGG-2-mediated xenophagy as playing a role in clearance, but they show it is not required. It is possible that LGG-2 does mediate clearance through xenophagy, and this process is redundant with another form of clearance (Figure 6). Given that *C. elegans* and *Nematocida* species appear to have a long co-evolutionary relationship,^17^ it is possible that *C. elegans* evolved additional forms of clearance as part of an arms race between this host and its natural pathogen. Studies in other systems have described pathogen virulence factors that suppress host autophagy, for example through removing LC3 conjugation from the autophagosomal membrane.^9^ Perhaps because another pathogen suppressed LGG-2-mediated autophagy in its evolutionary past, *C. elegans* evolved a separate pathway for clearance that functions here in the absence of LGG-2 to clear *N. ironsii*. Regardless of the reason, it is clear that HW animals have a pathway separate from LGG-2 that mediates clearance of *N. ironsii*. Given the extremely close correlation between ubiquitin targeting and *Nematocida* clearance (Figures 1 and 2), perhaps ubiquitin is functionally relevant for this pathway, and serves to recruit another type of degradative machinery to the parasite in parallel to LGG-2. Unfortunately, because ubiquitin is an essential gene and functions in many processes, it is difficult to test a functional role in clearance specifically. It is also possible that the ubiquitin targeting we see is not functionally relevant but is simply correlative with clearance. Indeed, there are several other studies where targeting of autophagic machinery to microbes is not indicative of a functional role in clearance.^10, 41^ It is difficult to thoroughly test a role for autophagy in the clearance of *Nematocida* from HW animals, because many autophagy proteins are essential, and as mentioned above, germline RNAi knock-down in HW is ineffective. Nonetheless, our findings suggest *C. elegans* evolved additional, non-xenophagic strategies for clearance of natural pathogens.

Interestingly, our results point to a clearance-independent role for *lgg-2* in controlling colonization of *Nematocida* inside of intestinal cells of HW animals, after the spores are ingested into the intestinal lumen. One explanation for these results is that *lgg-2* has a role in regulating the viability of microsporidia spores in the extracellular lumenal space, or in regulating their ability to fire the polar tube for invasion of intestinal cells. If so, this effect would be consistent with LGG-2 acting through secretion of anti-microbial compounds, as has been shown for regulation of lysozyme secretion by the autophagy pathway in the mammalian intestine.^12^ Another possibility is that *lgg-2* regulates the intracellular establishment of sporoplasms after they have been delivered into the cytoplasm by the polar tube, and leads to their degradation so quickly that we do not detect them in the FISH assay. Nothing is known about whether host cells control this initial stage of microsporidia invasion, as polar tube firing is described as a mechanical rupture that ‘forces’ a parasite cell into the host cell.^18^ However, it is exciting to consider that host cells may regulate this step via LGG-2/LC3. It is important to note that LC3 has non-autophagy roles in immunity, for example through LC3-associated phagocytosis or LAP.^42, 43^ During LAP, microbes that have been phagocytosed by host cells are targeted by LC3 and then degraded through phagosomal/lysosomal fusion. While we have not seen a separate membrane around *N. parisii* sporoplasms suggesting they are not phagocytosed,^20^ it is possible that LC3/LGG-2 is recruited to sporoplasms that are in direct contact with cytoplasm, and directs their immediate degradation through some process related to LAP.

Our observations that *lgg-2* controls sporoplasm levels at both 15 minutes post-inoculation and 3 hpi are arguably the earliest stages at which a host factor has been shown to control microsporidia load in any system. Studies of microsporidia species that infect mammalian cells have implicated host glycosaminoglycans^44^ and a transferrin receptor protein^45^ in regulation of early steps, but the exact stage and mechanism by which they act is unknown. It will be interesting to further explore this early role for *lgg-2* to shed light on the poorly understood question of how hosts control microsporidia infection.

## Acknowledgements

We thank the Malene Hansen Lab, Arshad Desai/Karen Oegema labs and Renaud Legouis lab for reagents. We thank Cheng-Ju Kuo for technical support. We thank Malene Hansen, Robert Luallen, Johan Panek, Kirthi Reddy, Ivana Sfarcic, Eillen Tecle, and Ryan Underwood for comments on the manuscript.

---

**Table S1.**
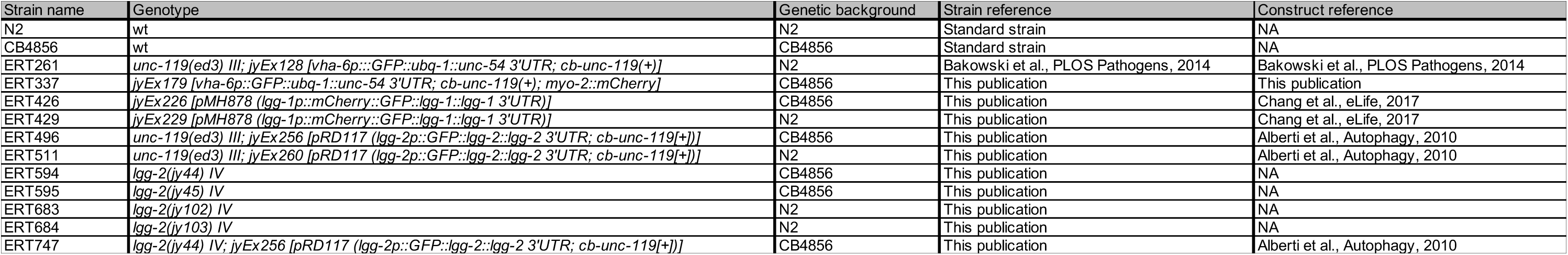
List of strains used in this study

**Table S2.**
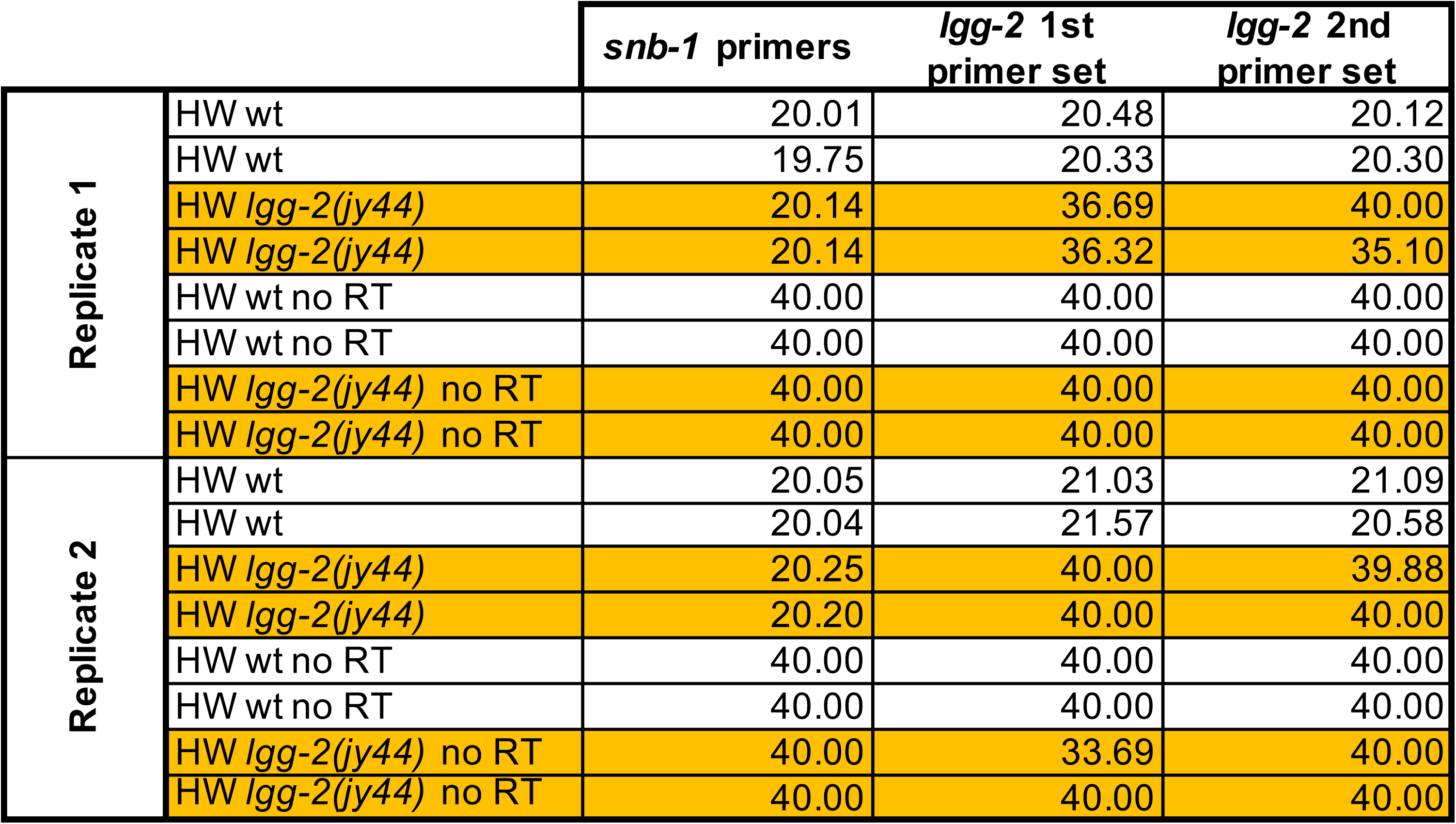
qRT-PCR measurements of *lgg-2* mRNA transcripts in HW wild-type and *lgg-2(jy44)* animals. Ct (cycle threshold) values for primer-template combinations are indicated in the table. The Ct is defined as the number of cycles required for the fluorescent signal to cross the threshold (ie exceeds background level), with higher Ct values indicating lower levels of starting template. A Ct value of 40 shows there was no signal detected for the *lgg-2* primers in the *lgg-2(jy44)* mutant. (Because there was no signal we did not provide a quantitative comparison among samples, which is the typical method for reporting qPCR data, e.g. ^46^). noRT samples represent negative controls in which reverse transcriptase was not added during cDNA synthesis.

**Table S3.**
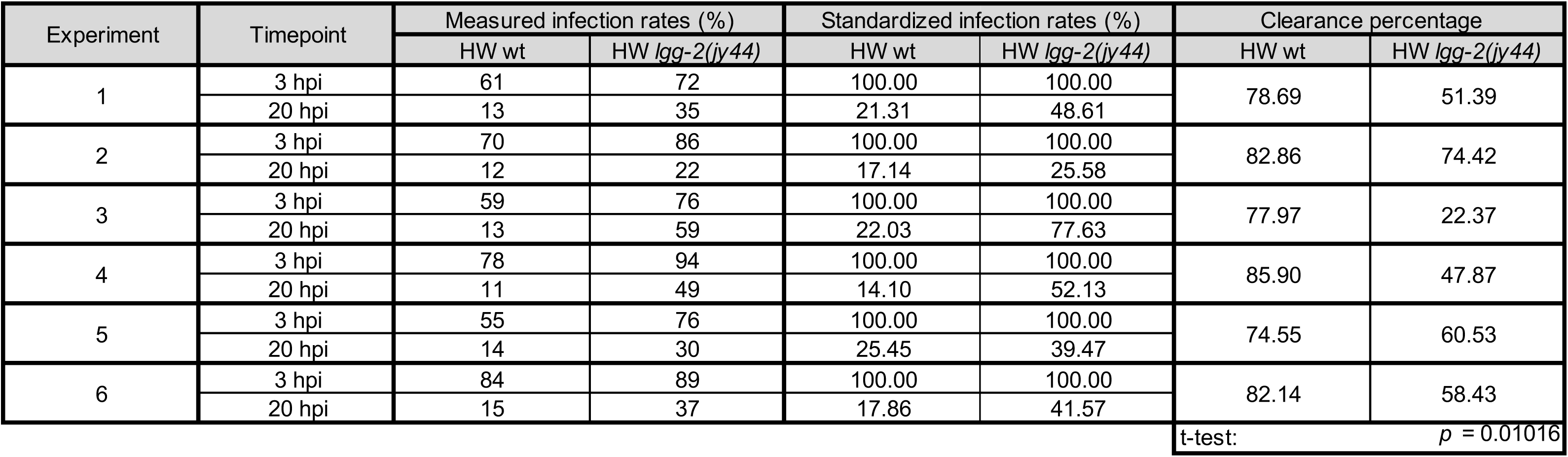
Statistical analysis of *N. ironsii* clearance in HW wt and HW *lgg-2(jy44)* animals at 20 hpi. To compare clearance efficiencies between HW wild-type and *lgg-2(jy44)* animals in six separate experiments, infection rates in each experiment at 3 hpi were standardized to a starting value of 100%. Then infection rates at 20 hpi were proportionally re-calculated and subtracted from 100% to determine the clearance percentage for each strain. Student’s t-test was used to compare clearance and calculate the *p* value.

**Figure S1.**
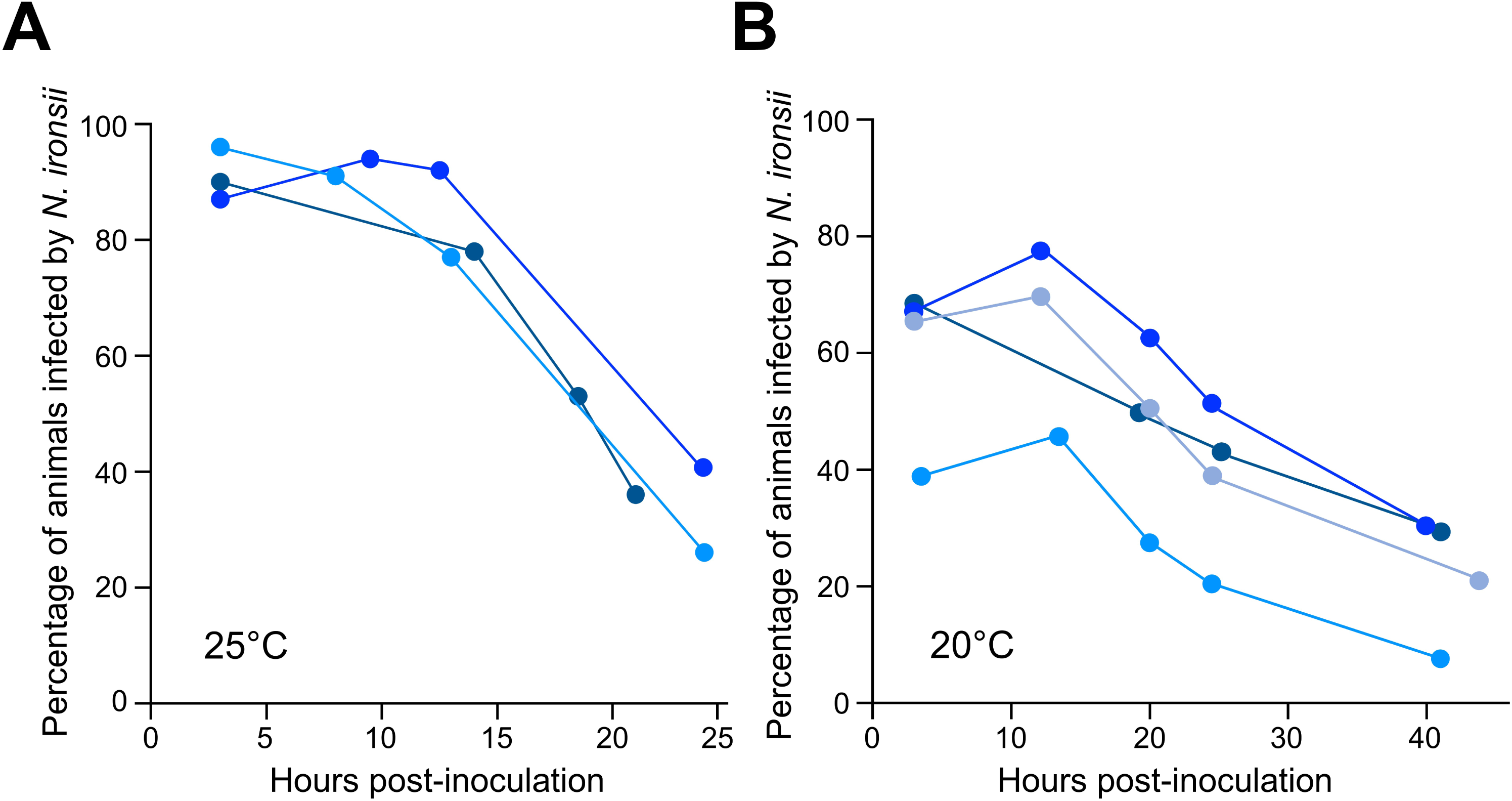
Clearance of *N. ironsii* infections occurs gradually over time in HW animals at both 25°C and 20°C. (A) HW L1 animals were pulse-inoculated with *N. ironsii* spores and fractions of the population were fixed at several time points afterwards to assess the frequency of infection. Experiments were carried out at 25°C. Data from three independent experiments are shown in blue lines. (B) Same experimental setup described above in (A), only the four experiments were carried out at 20°C.

**Figure S2.**
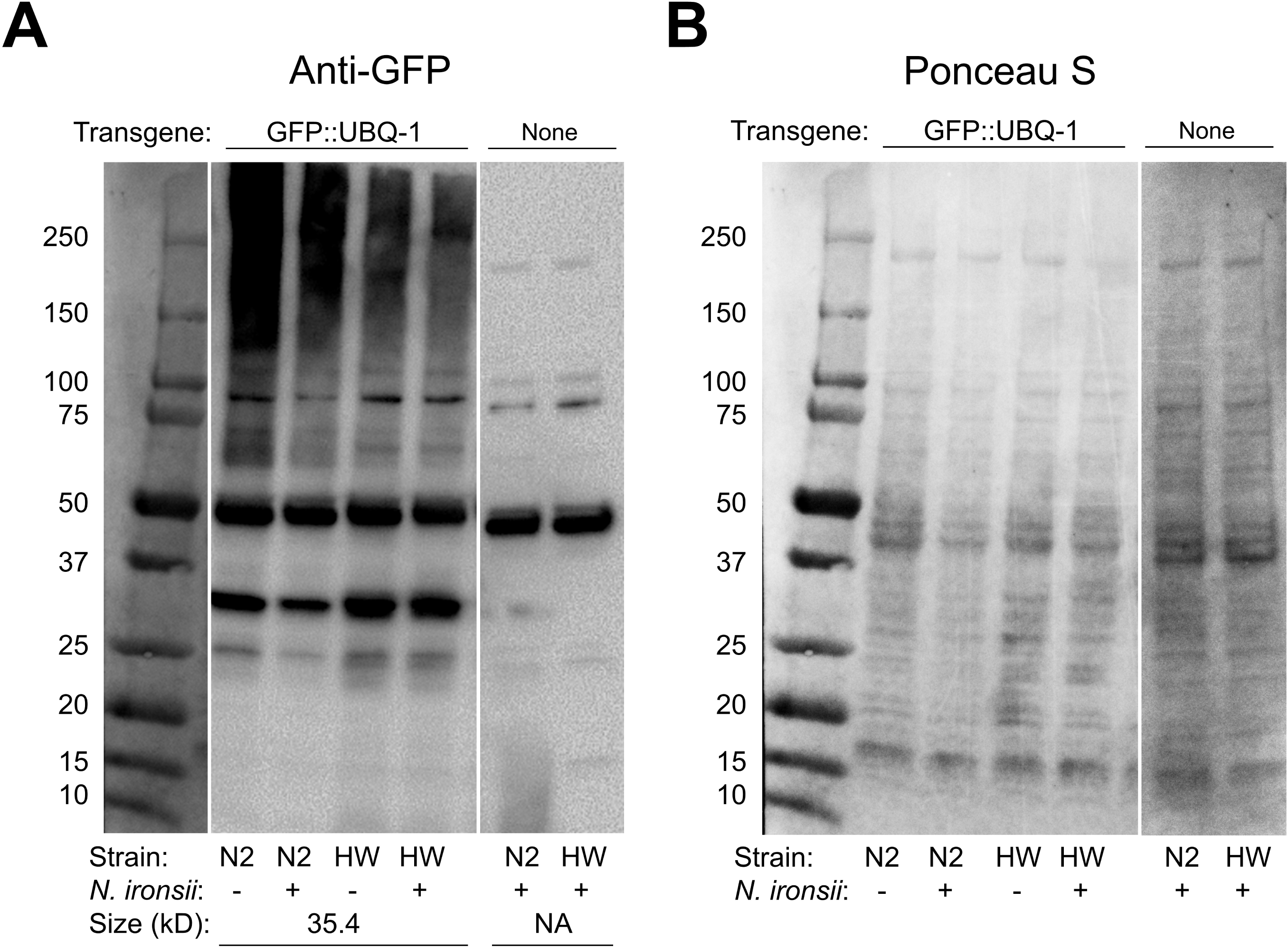
GFP::UBQ-1 fusion Western blot analysis in transgenic N2 and HW strains. (A) Proteins were extracted from uninfected and *N. ironsii* infected animals (15 hpi) and analyzed by SDS-PAGE followed by Western blot using an anti-GFP antibody. Background staining of proteins from non-transgenic infected animals is shown in the last two columns. The predicted size of GFP::UBQ-1 is 35.4 kD. (B) Loading of all proteins per sample visualized by Ponceau S staining.

**Figure S3.**
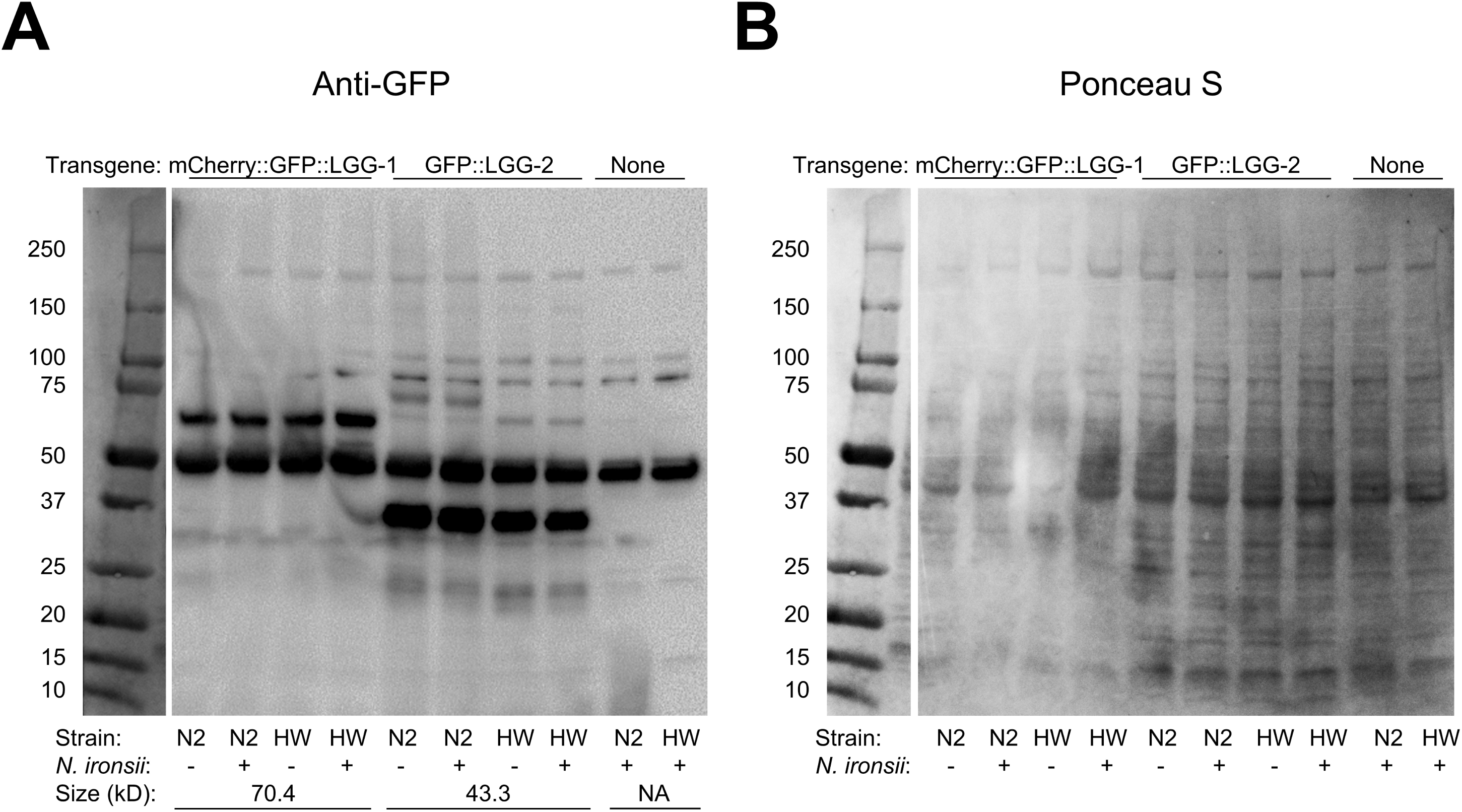
GFP::LGG-1 and GFP::LGG-2 fusion Western blot analysis in transgenic N2 and HW strains. (A) Proteins were extracted from uninfected and *N. ironsii* infected animals (15 hpi) and analyzed by SDS-PAGE followed by Western blot using an anti-GFP antibody. Background staining of proteins from non-transgenic infected animals is shown in the last two columns. The predicted size of GFP::LGG-1 is 70.4 kD and the predicted size of GFP::LGG-2 is 43.3 kD. (B) Loading of all proteins per sample visualized by Ponceau S staining.

**Figure S4.**
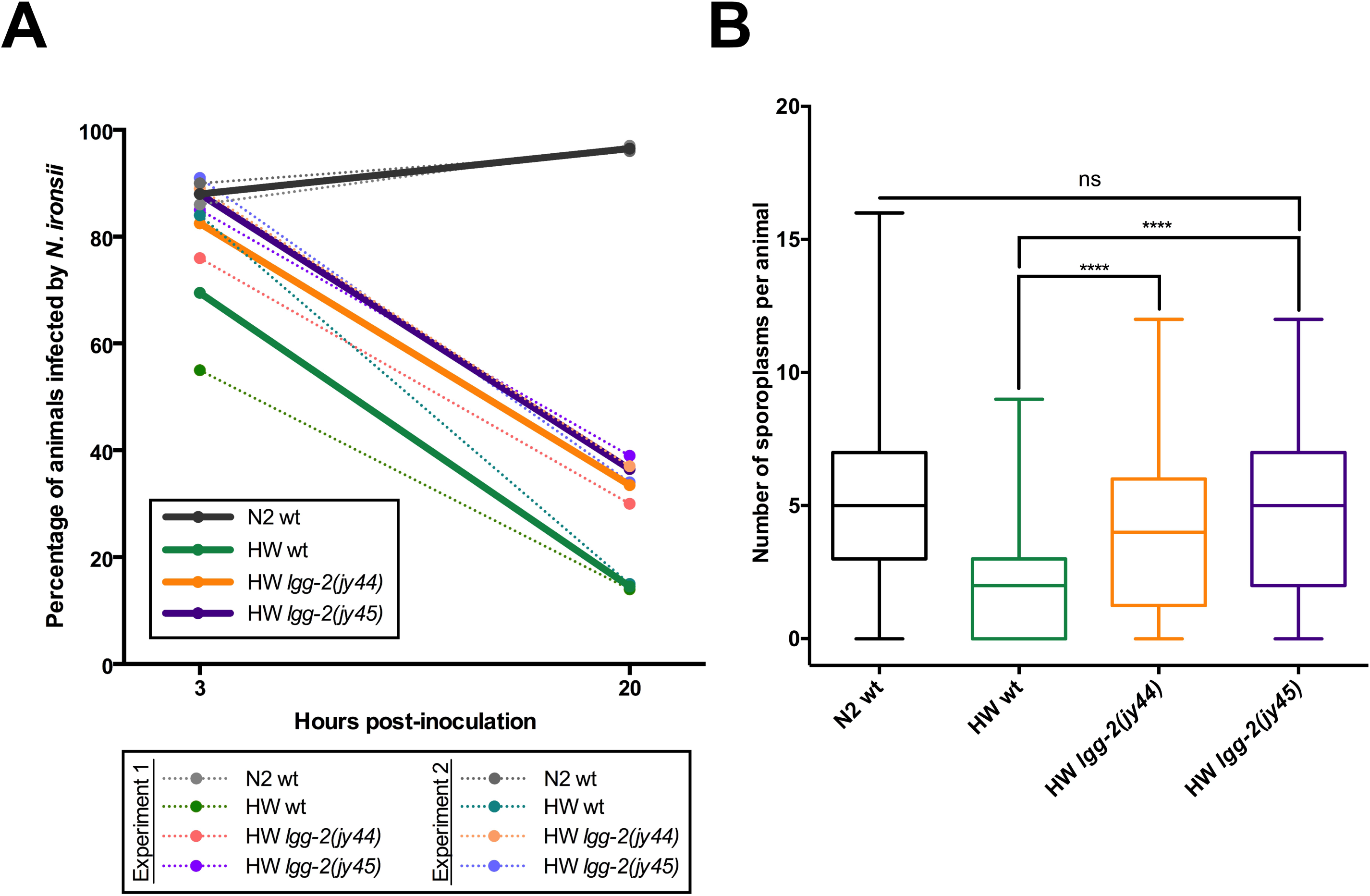
Different *lgg-2* deletion mutants in the HW background exhibit similar phenotypes. (A) *N. ironsii* clearance in *lgg-2*(*jy44)* mutants (orange line) and *lgg-2(jy45)* mutants (purple line). Thick lines represent average values from two independent experiments; each replicate is shown with a dotted line. HW and N2 wild-type controls are shown in green and gray lines, respectively. (B) Box-and-whiskers plot shows similar *N. ironsii* infection rate between *lgg-2*(*jy44)* and *lgg-2(jy45)* alleles (3 hpi). Each box represents 50% of the data from two independent experiments closest to the median value (line in the box). Whiskers span the values outside of the box. A student’s t-test was used to calculate *p* values; *p* < 0.001 is indicated with four asterisks; ns indicates non-significant difference (*p* > 0.05). (A, B) All experiments were performed at 25°C.

**Figure S5.**
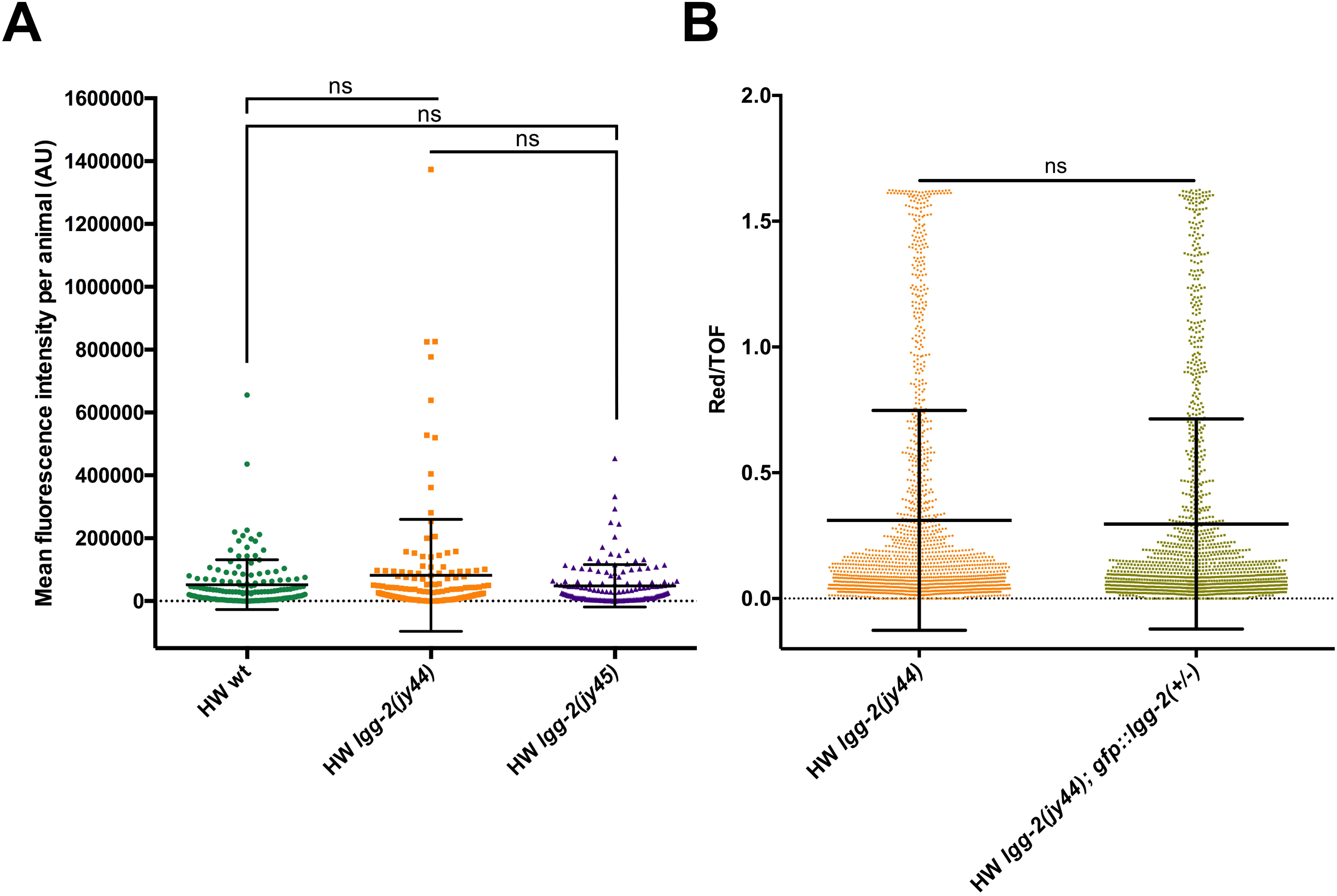
Bead feeding assays indicate similar feeding rates between analyzed strains. (A, B) Fluorescent bead accumulation in the intestines of infected L1 animals after 30 minutes. (A) *lgg2(jy44)* and *lgg-2(jy45)* mutants show similar feeding rate to HW wild-type animals. Feeding assay was performed in triplicate, 50 animals were analyzed per strain per experiment. Average red fluorescence intensities per whole animal are shown in arbitrary units (AU) on y-axis. Each dot represents one animal. (B) Mixed population of GFP::LGG-2 rescued and non-rescued *lgg-2(jy44)* animals have a similar feeding rate as *lgg-2(jy44)* strain. Fluorescence levels measurements were standardized to the body length for each animal. More than 1350 animals were analyzed from three individual experiments combined. (A, B) A student’s t-test was used to calculate *p* values; ns indicates non-significant difference (*p* > 0.05). All experiments were performed at 25°C.

**Figure S6.**
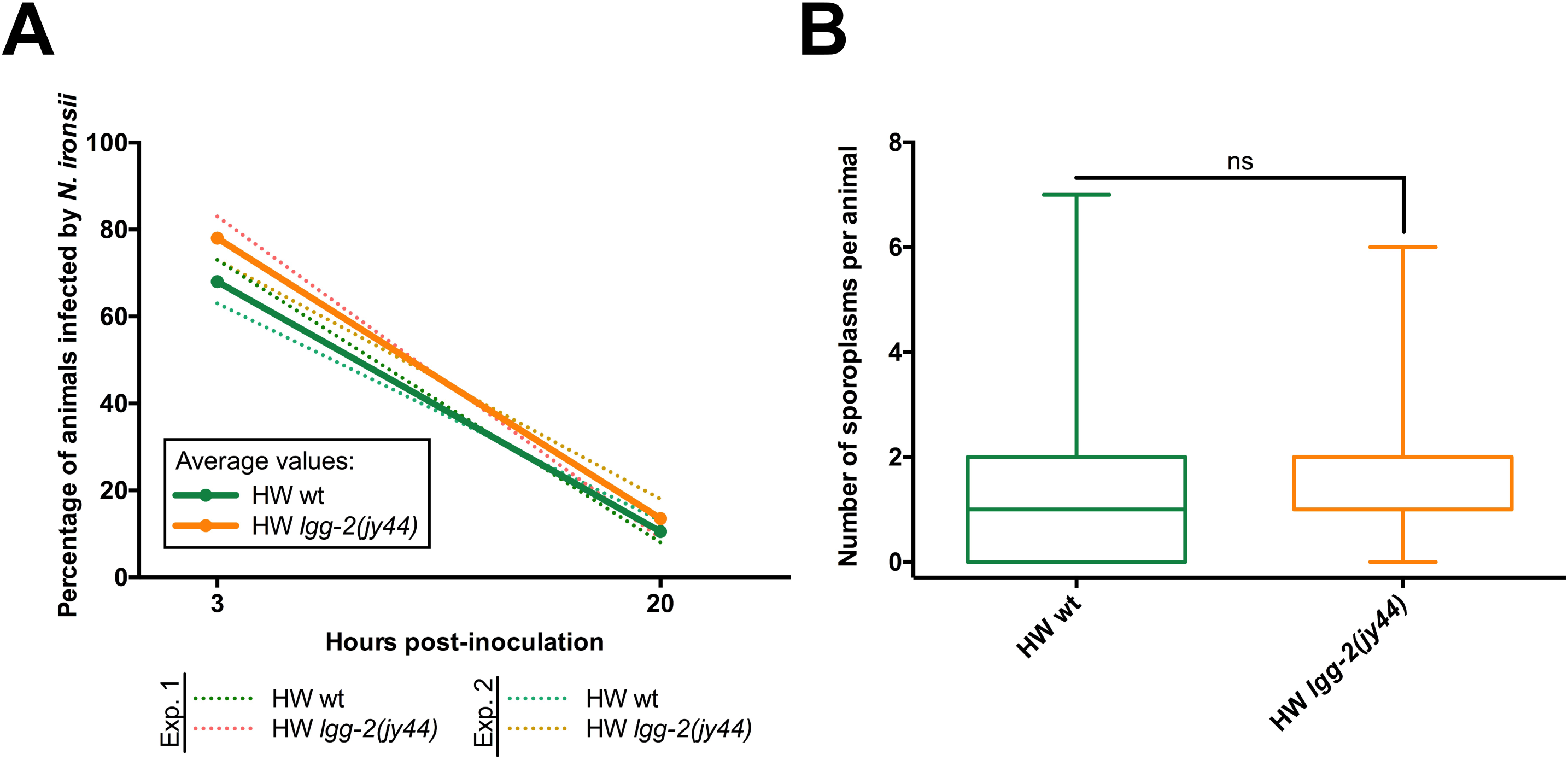
Ability to clear infection is similar in wild-type HW animals and *lgg-2* deletion mutants if initial infection rate is similar. (A, B) *N. ironsii* clearance in HW wild-type and *lgg-2(jy44)* animals is similar if HW wt animals are infected with a higher dose of microsporidia spores than HW *lgg-2(jy44)* mutants, to achieve similar initial intestinal colonization in both strains. (A) *N. ironsii* clearance in HW wild-type animals (green lines) and *lgg-2*(*jy44)* mutants (orange lines). Thick lines represent average values of two experiments; dotted lines indicate results from individual experiments. 100 animals were analyzed per strain at 3 hpi and 20 hpi (x-axis). (B) Box-and-whiskers plot shows similar *N. ironsii* infection rate between HW wild-type and *lgg-2(jy44)* mutant animals. Each box represents 50% of the data closest to the median value (line in the box). Note that the median value for HW *lgg-2(jy44)* sample is two sporoplasms per animal and that it overlaps with the upper boundary of the box. Whiskers span the values outside of the box. A student’s t-test was used to calculate *p* values; ns indicates non-significant difference (*p* > 0.05).

**Figure S7.**
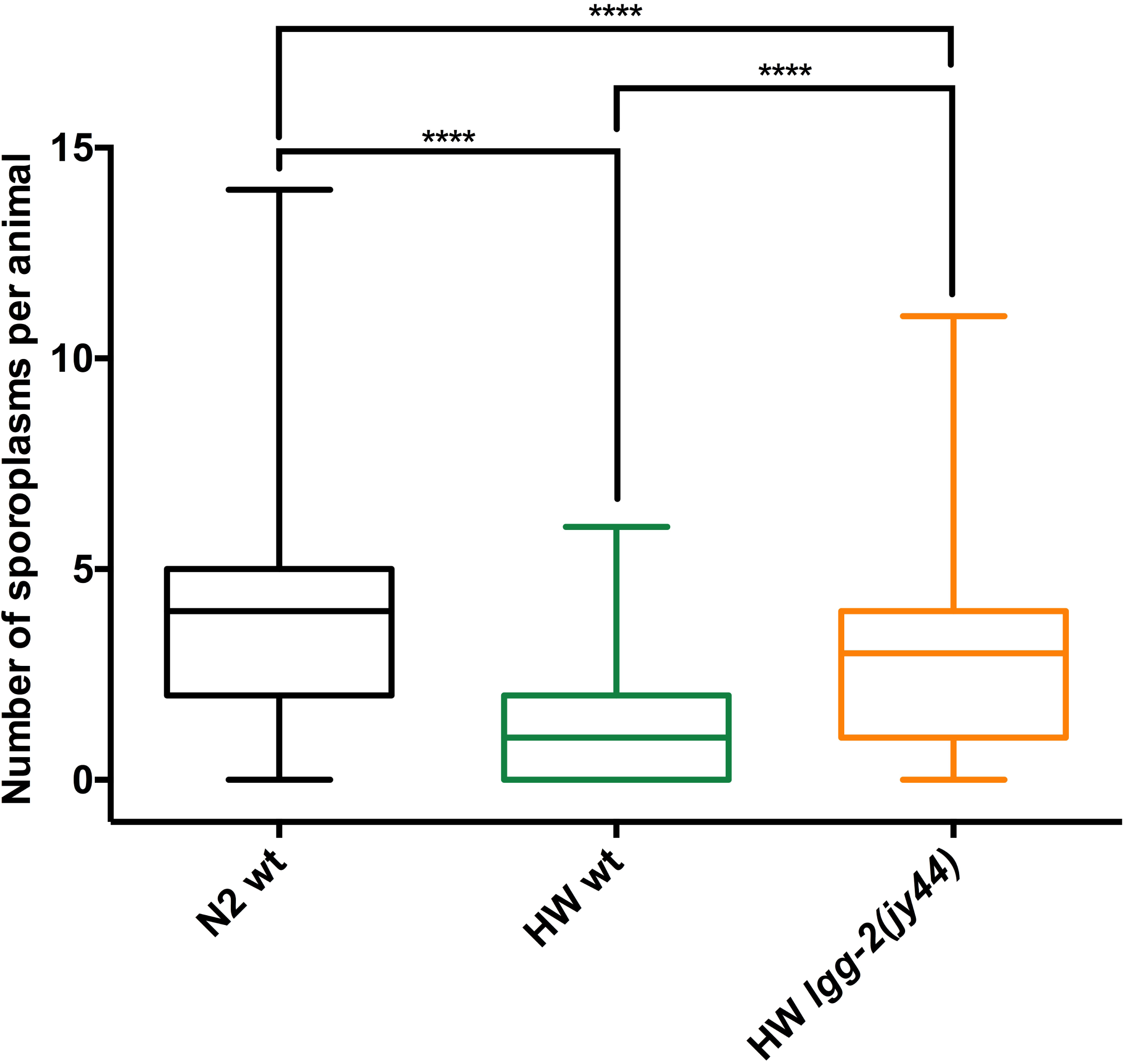
Short-term exposure to high dose of pathogen causes higher infection rate in HW *lgg-2* deletion mutants. Significant difference in pathogen load is observed between HW wild-type and HW *lgg-2* deletion mutant animals after 15 minutes incubation with 12 million *N. ironsii* spores (6 times more spores than in other clearance assays with *lgg-2* mutants). Results from two independent experiments are shown as box-and-whisker plots, indicating the number of *N. ironsii* sporoplasms per animal at 3 hpi. Each box represents 50% of the data closest to the median value (line in the box). Whiskers span the values outside of the box. 400 animals were analyzed for each strain. A student’s t-test was used to calculate *p* values; *p* < 0.001 is indicated with four asterisks; ns indicates non-significant difference (*p* > 0.05). Experiments were performed in liquid culture at 25°C.

**Figure S8.**
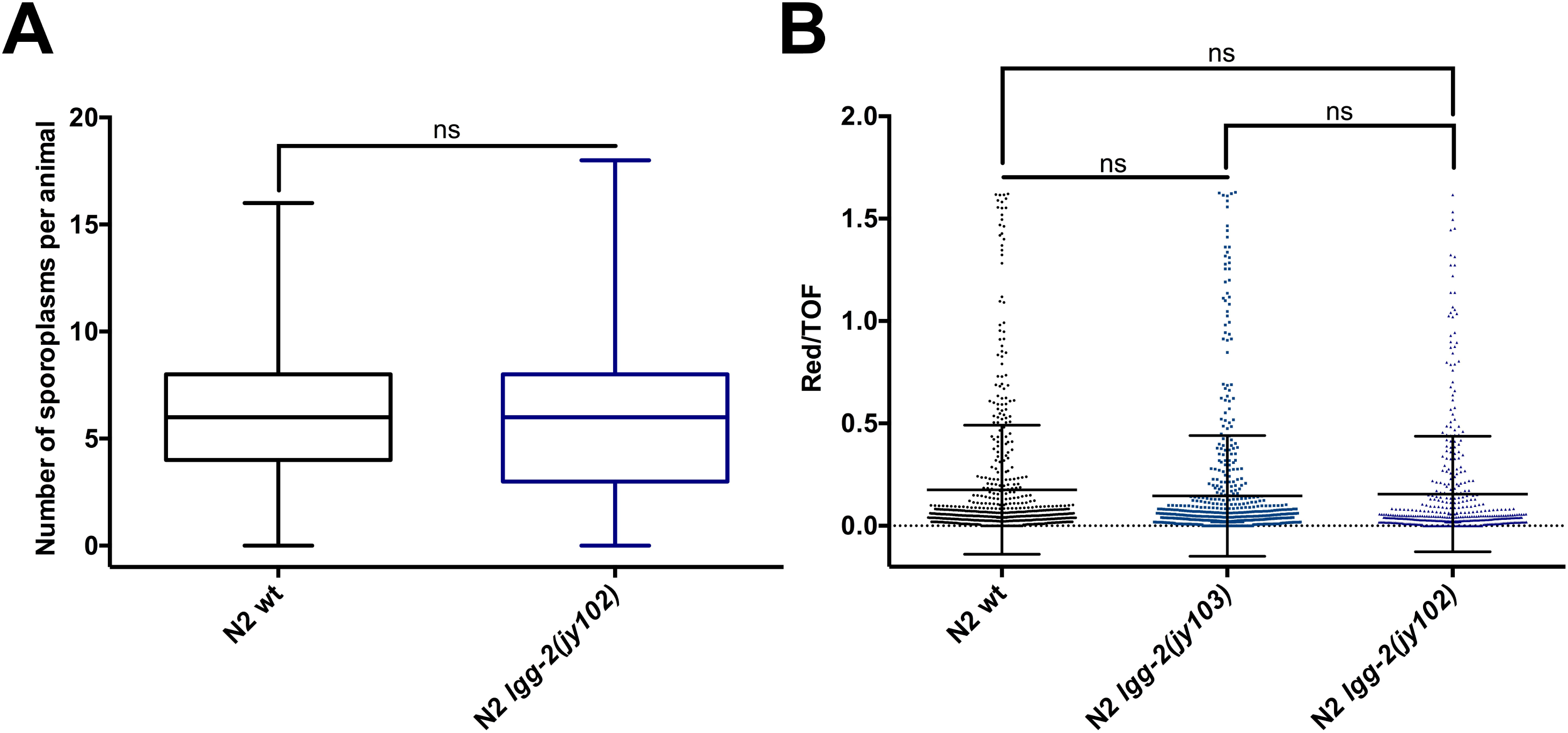
Deletion of *lgg-2* in the N2 background does not affect *N. ironsii* colonization. (A) *lgg-2(jy102)* mutants show similar infection rate as the wild type control. Results from two independent experiments are shown as box-and-whisker plots, indicating the number of *N. ironsii* sporoplasms per animal at 3 hpi. Each box represents 50% of the data closest to the median value (line in the box). Whiskers span the values outside of the box. 200 animals were analyzed for each strain. (B) N2 wild type and two N2 *lgg-2* mutant strains (*lgg-2(jy103)* and *lgg-2(102)*) have similar feeding rate. Bead fluorescence is standardized to TOF. At least 450 animals were analyzed for each strain, combined from three replicate experiments. (A, B) A student’s t-test was used to calculate *p* values; ns indicates non-significant difference (*p* > 0.05). Experiments were performed at 25°C.

## References

1. Wang L, Pittman KJ, Barker JR, Salinas RE, Stanaway IB, Williams GD, et al. An Atlas of Genetic Variation Linking Pathogen-Induced Cellular Traits to Human Disease. Cell host & microbe 2018; 24:308–23 e6.

2. Rioux JD, Xavier RJ, Taylor KD, Silverberg MS, Goyette P, Huett A, et al. Genome-wide association study identifies new susceptibility loci for Crohn disease and implicates autophagy in disease pathogenesis. Nature genetics 2007; 39:596–604.

3. Khor B, Gardet A, Xavier RJ. Genetics and pathogenesis of inflammatory bowel disease. Nature 2011; 474:307–17.

4. Lassen KG, Xavier RJ. Genetic control of autophagy underlies pathogenesis of inflammatory bowel disease. Mucosal Immunol 2017; 10:589–97.

5. Benjamin JL, Sumpter R, Jr., Levine B, Hooper LV. Intestinal epithelial autophagy is essential for host defense against invasive bacteria. Cell host & microbe 2013; 13:723–34.

6. Yin Z, Pascual C, Klionsky DJ. Autophagy: machinery and regulation. Microb Cell 2016; 3:588–96.

7. Randow F, Youle RJ. Self and nonself: how autophagy targets mitochondria and bacteria. Cell host & microbe 2014; 15:403–11.

8. Miller C, Celli J. Avoidance and Subversion of Eukaryotic Homeostatic Autophagy Mechanisms by Bacterial Pathogens. J Mol Biol 2016; 428:3387–98.

9. McEwan DG. Host-pathogen interactions and subversion of autophagy. Essays Biochem 2017; 61:687–97.

10. Mitchell G, Isberg RR. Innate Immunity to Intracellular Pathogens: Balancing Microbial Elimination and Inflammation. Cell host & microbe 2017; 22:166–75.

11. Ktistakis NT, Tooze SA. Digesting the Expanding Mechanisms of Autophagy. Trends in cell biology 2016; 26:624–35.

12. Bel S, Pendse M, Wang Y, Li Y, Ruhn KA, Hassell B, et al. Paneth cells secrete lysozyme via secretory autophagy during bacterial infection of the intestine. Science (New York, NY 2017; 357:1047–52.

13. Kuo CJ, Hansen M, Troemel E. Autophagy and innate immunity: Insights from invertebrate model organisms. Autophagy 2018; 14:233–42.

14. Chen Y, Scarcelli V, Legouis R. Approaches for Studying Autophagy in Caenorhabditis elegans. Cells 2017; 6.

15. Zhang H, Chang JT, Guo B, Hansen M, Jia K, Kovacs AL, et al. Guidelines for monitoring autophagy in Caenorhabditis elegans. Autophagy 2015; 11:9–27.

16. Frezal L, Felix MA. C. elegans outside the Petri dish. Elife 2015; 4.

17. Zhang G, Sachse M, Prevost MC, Luallen RJ, Troemel ER, Felix MA. A Large Collection of Novel Nematode-Infecting Microsporidia and Their Diverse Interactions with Caenorhabditis elegans and Other Related Nematodes. PLoS pathogens 2016; 12:e1006093.

18. Vavra J, Lukes J. Microsporidia and ‘the art of living together’. Advances in parasitology 2013; 82:253–319.

19. Troemel ER, Felix MA, Whiteman NK, Barriere A, Ausubel FM. Microsporidia are natural intracellular parasites of the nematode *Caenorhabditis elegans*. PLoS biology 2008; 6:2736–52.

20. Szumowski SC, Botts MR, Popovich JJ, Smelkinson MG, Troemel ER. The small GTPase RAB-11 directs polarized exocytosis of the intracellular pathogen N. parisii for fecal-oral transmission from C. elegans. Proceedings of the National Academy of Sciences of the United States of America 2014; 111:8215–20.

21. Bakowski MA, Desjardins CA, Smelkinson MG, Dunbar TA, Lopez-Moyado IF, Rifkin SA, et al. Ubiquitin-mediated response to microsporidia and virus infection in C. elegans. PLoS pathogens 2014; 10:e1004200.

22. Gomez-Diaz C, Ikeda F. Roles of ubiquitin in autophagy and cell death. Semin Cell Dev Biol 2018.

23. Jia K, Thomas C, Akbar M, Sun Q, Adams-Huet B, Gilpin C, et al. Autophagy genes protect against Salmonella typhimurium infection and mediate insulin signaling-regulated pathogen resistance. Proceedings of the National Academy of Sciences of the United States of America 2009; 106:14564–9.

24. Visvikis O, Ihuegbu N, Labed SA, Luhachack LG, Alves AM, Wollenberg AC, et al. Innate host defense requires TFEB-mediated transcription of cytoprotective and antimicrobial genes. Immunity 2014; 40:896–909.

25. Zou CG, Ma YC, Dai LL, Zhang KQ. Autophagy protects C. elegans against necrosis during Pseudomonas aeruginosa infection. Proceedings of the National Academy of Sciences of the United States of America 2014; 111:12480–5.

26. Kirienko NV, Ausubel FM, Ruvkun G. Mitophagy confers resistance to siderophore-mediated killing by Pseudomonas aeruginosa. Proceedings of the National Academy of Sciences of the United States of America 2015; 112:1821–6.

27. Balla KM, Andersen EC, Kruglyak L, Troemel ER. A wild C. elegans strain has enhanced epithelial immunity to a natural microsporidian parasite. PLoS pathogens 2015; 11:e1004583.

28. Reinke AW, Balla KM, Bennett EJ, Troemel ER. Identification of microsporidia host-exposed proteins reveals a repertoire of rapidly evolving proteins. Nat Commun 2017; 8:14023.

29. Cohen LB, Troemel ER. Microbial pathogenesis and host defense in the nematode C. elegans. Current opinion in microbiology 2015; 23:94–101.

30. Irazoqui JE, Urbach JM, Ausubel FM. Evolution of host innate defence: insights from Caenorhabditis elegans and primitive invertebrates. Nature reviews 2010; 10:47–58.

31. Brenner S. The genetics of *Caenorhabditis elegans*. Genetics 1974; 77:71–94.

32. Chang JT, Kumsta C, Hellman AB, Adams LM, Hansen M. Spatiotemporal regulation of autophagy during Caenorhabditis elegans aging. Elife 2017; 6.

33. Manil-Segalen M, Lefebvre C, Jenzer C, Trichet M, Boulogne C, Satiat-Jeunemaitre B, et al. The C. elegans LC3 acts downstream of GABARAP to degrade autophagosomes by interacting with the HOPS subunit VPS39. Dev Cell 2014; 28:43–55.

34. Alberti A, Michelet X, Djeddi A, Legouis R. The autophagosomal protein LGG-2 acts synergistically with LGG-1 in dauer formation and longevity in C. elegans. Autophagy 2010; 6:622–33.

35. Cuomo CA, Desjardins CA, Bakowski MA, Goldberg J, Ma AT, Becnel JJ, et al. Microsporidian genome analysis reveals evolutionary strategies for obligate intracellular growth. Genome Res 2012; 22:2478–88.

36. Estes KA, Szumowski SC, Troemel ER. Non-lytic, actin-based exit of intracellular parasites from *C. elegans* intestinal cells. PLoS pathogens 2011; 7:e1002227.

37. Paix A, Folkmann A, Rasoloson D, Seydoux G. High Efficiency, Homology-Directed Genome Editing in Caenorhabditis elegans Using CRISPR-Cas9 Ribonucleoprotein Complexes. Genetics 2015; 201:47–54.

38. Nguyen TN, Padman BS, Usher J, Oorschot V, Ramm G, Lazarou M. Atg8 family LC3/GABARAP proteins are crucial for autophagosome-lysosome fusion but not autophagosome formation during PINK1/Parkin mitophagy and starvation. The Journal of cell biology 2016; 215:857–74.

39. Pollard DA, Rockman MV. Resistance to germline RNA interference in a Caenorhabditis elegans wild isolate exhibits complexity and nonadditivity. G3 (Bethesda) 2013; 3:941–7.

40. Tijsterman M, Okihara KL, Thijssen K, Plasterk RH. PPW-1, a PAZ/PIWI protein required for efficient germline RNAi, is defective in a natural isolate of C. elegans. Curr Biol 2002; 12:1535–40.

41. Mitchell G, Cheng MI, Chen C, Nguyen BN, Whiteley AT, Kianian S, et al. Listeria monocytogenes triggers noncanonical autophagy upon phagocytosis, but avoids subsequent growth-restricting xenophagy. Proceedings of the National Academy of Sciences of the United States of America 2018; 115:E210–E7.

42. Schille S, Crauwels P, Bohn R, Bagola K, Walther P, van Zandbergen G. LC3-associated phagocytosis in microbial pathogenesis. Int J Med Microbiol 2017.

43. Jenzer C, Simionato E, Largeau C, Scarcelli V, Lefebvre C, Legouis R. Autophagy mediates phosphatidylserine exposure and phagosome degradation during apoptosis through specific functions of GABARAP/LGG-1 and LC3/LGG-2. Autophagy 2018:1–14.

44. Hayman JR, Southern TR, Nash TE. Role of sulfated glycans in adherence of the microsporidian Encephalitozoon intestinalis to host cells in vitro. Infection and immunity 2005; 73:841–8.

45. Han B, Polonais V, Sugi T, Yakubu R, Takvorian PM, Cali A, et al. The role of microsporidian polar tube protein 4 (PTP4) in host cell infection. PLoS pathogens 2017; 13:e1006341.

46. Pfaffl MW. A new mathematical model for relative quantification in real-time RT-PCR. Nucleic Acids Res 2001; 29:e45.

